# Fxr signaling and microbial metabolism of bile salts in the zebrafish intestine

**DOI:** 10.1101/2020.12.13.422569

**Authors:** Jia Wen, Gilberto Padilla Mercado, Alyssa Volland, Heidi L. Doden, Colin R. Lickwar, Taylor Crooks, Genta Kakiyama, Cecelia Kelly, Jordan L. Cocchiaro, Jason M. Ridlon, John F. Rawls

## Abstract

Bile salt synthesis, secretion into the intestinal lumen, and resorption in the ileum occurs in all vertebrate classes. In mammals, bile salt composition is determined by host and microbial enzymes, affecting signaling through the bile salt-binding transcription factor Farnesoid X receptor (Fxr). However, these processes in other vertebrate classes remain poorly understood. We show that key components of hepatic bile salt synthesis and ileal transport pathways are conserved and under control of Fxr in zebrafish. Zebrafish bile salts consist primarily of a C_27_ bile alcohol and a C_24_ bile acid which undergo multiple microbial modifications including bile acid deconjugation that augments Fxr activity. Using single-cell RNA sequencing, we provide a cellular atlas of the zebrafish intestinal epithelium and uncover roles for Fxr in transcriptional and differentiation programs in ileal and other cell types. These results establish zebrafish as a non-mammalian vertebrate model for studying bile salt metabolism and Fxr signaling.

## Introduction

Bile salts are the end product of cholesterol catabolism in the liver of all vertebrates (*1*). Upon lipid ingestion, bile salts are released into the duodenum as emulsifiers to solubilize lipids and are then reabsorbed by the ileum into the portal vein to return to the liver, a process known as enterohepatic circulation. Bile salts also act as signaling molecules that exert diverse effects by activating nuclear or membrane-bound receptors (*2*). This includes the nuclear receptor Farnesoid X receptor (FXR/NR1H4), an evolutionarily conserved transcription factor that uses bile salts as endogenous ligands (*3*). Upon binding with bile salts, FXR regulates a large number of target genes involved in bile salt, lipid, and glucose metabolism (*4*). FXR activity can be modulated by the chemical structure of bile salts, which differ considerably across vertebrate species (*5*). For example, fish and amphibians contain predominantly 27-carbon (C_27_) bile alcohols, whereas mammals mainly possess 24-carbon (C_24_) bile acids (*1*). Even within the same species, there can be substantial diversity in bile salt structures. One key contributor to this diversity is the gut microbiota, which can modify the side chain(s) or stereostructure of the conjugated primary bile salts synthesized by the liver (*6*). This leads to the production of various unconjugated or secondary bile salts in the intestine with different activities towards FXR, therefore altering FXR-mediated signaling pathways. Though bile salts and FXR are present in diverse vertebrate species (*7, 8*), our knowledge about bile salt-FXR signaling has been almost entirely limited to humans and rodents. It remains unclear when this signaling axis arose and whether its functions changed over the course of vertebrate evolution. Further, despite mice in particular have been effective at revealing FXR functions, there are substantial differences between mice and humans, including bile salt composition, the effects of bile salt on Fxr, and Fxr-mediated metabolic activities (*9, 10*). Therefore, additional vertebrate models are needed to provide complementary perspectives into the mechanistic relationships between microbiota, bile salts, and FXR signaling, and to potentially reveal new functions of FXR.

The zebrafish (*Danio rerio*) has emerged as a powerful model for studying bile salt-related liver diseases due to their conserved mechanisms of liver and intestinal development and bile secretion, facile genetic and transgenic manipulations, and ease of monitoring host-microbiota interactions and other physiological processes *in vivo* (*11–14*). The genome of zebrafish possesses orthologs of many mammalian genes known to be involved in bile salt homeostasis, including bile salt transporters, bile salt synthesis enzymes, and FXR (*7, 12, 15, 16*). Further, genes involved in bile salt absorption are expressed in a conserved ileal region of the zebrafish intestine (*17*). However, the requirement for those zebrafish genes in enterohepatic circulation and bile salt signaling remains largely untested. Additionally, although primary bile salt composition in zebrafish has been assessed (*18, 19*), microbial metabolism of zebrafish bile salts has not been explored.

Here, we establish zebrafish as a non-mammalian vertebrate model to study the bile salt-Fxr signaling axis. We establish the evolutionary conservation of key components of this axis between zebrafish and mammals, and assess the contribution of zebrafish gut microbes to the modulation of the bile salt-Fxr signaling. Further, we uncover the requirements of zebrafish Fxr in gene expression and differentiation in multiple intestinal epithelial cell types using single-cell transcriptomics.

## Results

### Key components of the Fxr signaling pathway are conserved in zebrafish

We used CRISPR-Cas9 to generate *fxr* mutant zebrafish (*fxr*^-10/-10^, designated as *fxr*^-/-^) and then investigated the impacts on predicted Fxr targets (Fig 1A, S1A-B). *Fatty acid binding protein 6* (*fabp6*), the gene encoding the ileal bile acid binding protein, is a known Fxr target in mammals and is highly expressed in the zebrafish ileum (*17, 20*). Using a new reporter line *Tg(- 1.7fabp6:GFP)* that expresses GFP in the ileal epithelium under control of the 1.7kb *fabp6* promoter, we observed striking attenuation of GFP fluorescence in *fxr^-/-^* zebrafish compared to *fxr*^+/+^ wild-type (wt) controls (Fig 1B). This suggested that expression of *fabp6* in the ileum is dependent on Fxr in zebrafish as it is in mammals (*20*), and that this reporter line can be used to monitor Fxr activity *in vivo*. Examination of a larger panel of predicted Fxr target genes involved in bile salt homeostasis revealed similar transcriptional changes in *fxr^-/-^* zebrafish as seen in Fxr knockout mice (*12, 20–22*). This includes reduced expression of *fabp6*, the fibroblast growth factor *fgf19*, and the bile salt export pump *abcb11b*, along with induction of the *cyp7a1* which encodes the rate-limiting enzyme cholesterol 7alpha-hydroxylase in hepatic bile salt synthesis. Interestingly, the apical sodium-dependent bile acid transporter *slc10a2*, which is indirectly repressed by FXR in mice and humans, appeared to be positively regulated by Fxr in zebrafish, as *slc10a2* expression was reduced in *fxr*^-/-^ zebrafish. Nonetheless, these data reveal that Fxr is critical for the coordinated expression of bile salt metabolism genes in zebrafish as in mammals (Fig 1C).

**Figure 1.**
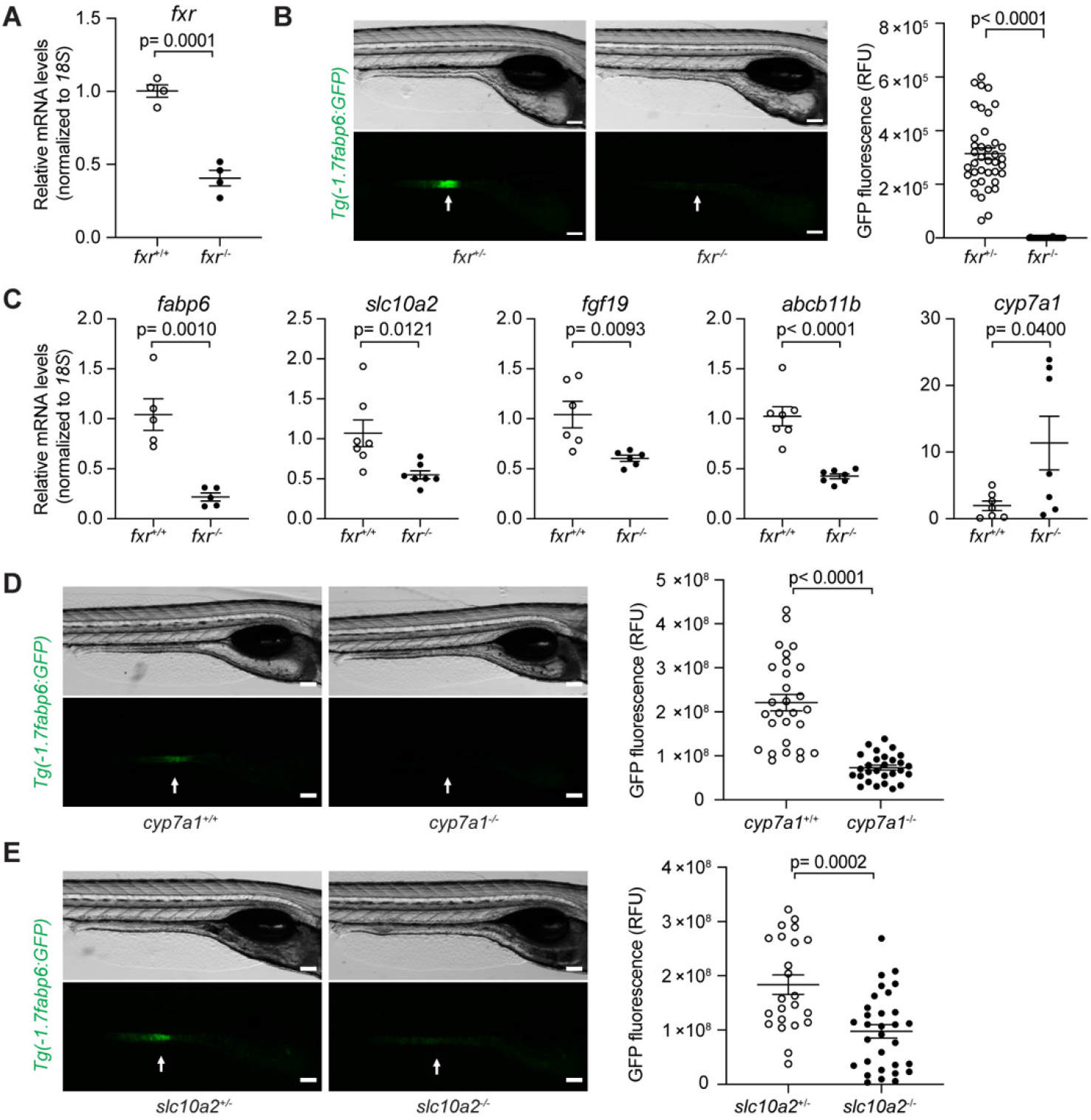
Genetic analyses reveal conserved key components of the bile salt-Fxr signaling axis in zebrafish. **A.** qRT-PCR comparing *fxr* expression in whole 7 dpf *fxr* wild type (*fxr*^+/+^) or *fxr* homozygous mutant (*fxr* ^-/-^) zebrafish larvae. **B.** Imaging and quantification of GFP fluorescence of the ileal region of 7 dpf *fxr*^+/+^ and *fxr*^-/-^ *Tg(-1.7fabp6:GFP)* larvae. The ileal region is indicated by arrows. **C.** qRT-PCR comparing expression of genes related to bile salt metabolism in dissected digestive tracts (including intestine, liver, pancreas, and gall bladder) of 7 dpf *fxr*^+/+^ or *fxr*^-/-^ larvae (for *fabp6*, *slc10a2*, and *fgf19*) or dissected liver of gender and size-matched adult *fxr*^+/+^ or *fxr*^-/-^ zebrafish (for *abcb11b* and *cyp7a1*). The results are represented as relative expression levels normalized *18S* (Mean±SEM). **D** and **E.** Imaging and quantification of GFP fluorescence of the ileal region of 7dpf *Tg(-1.7fabp6:GFP) cyp7a1*^+/+^ and *cyp7a1*^-/-^ larvae (D) or *slc10a2*^+/-^ and *slc10a2*^-/-^ larvae (E). The zebrafish ileal region is indicated by arrows. Scale bar = 100 μm in (B, D, E). Statistical significance was calculated by unpaired t-test. Shown are representative data from at least three independent experiments.

### Bile salt-mediated Fxr activation is conserved in zebrafish as in mammals

We next sought to test if bile salt mediated regulation of Fxr activity is conserved in zebrafish as in mammals. To do so, we first defined the level and diversity of zebrafish bile salts by analyzing the biliary bile extracted from pooled adult zebrafish gallbladders using ESI-LC/MS. Based on the mass ion, the major component (83.4%) of the purified zebrafish bile was determined to be 5α-cyprinol sulfate (5αCS), a C_27_ bile alcohol species commonly present in fishes (Fig. 2A) (*23, 24*). This was further validated by examining the 1H, 13C, COSY, and HSQC NMR spectra of this compound (Fig S2A). We also identified several minor bile salt species, including 8.8% taurocholic acid (TCA), a C_24_ bile acid commonly found in mammals, 7.8% 5α-cholestane- 3α,7α, 12α,26-tetrol sulfate, a precursor of 5αCS (*1, 23*), and a trace amount of the dehydrogenated form of 5αCS (Fig 2A).

**Figure 2.**
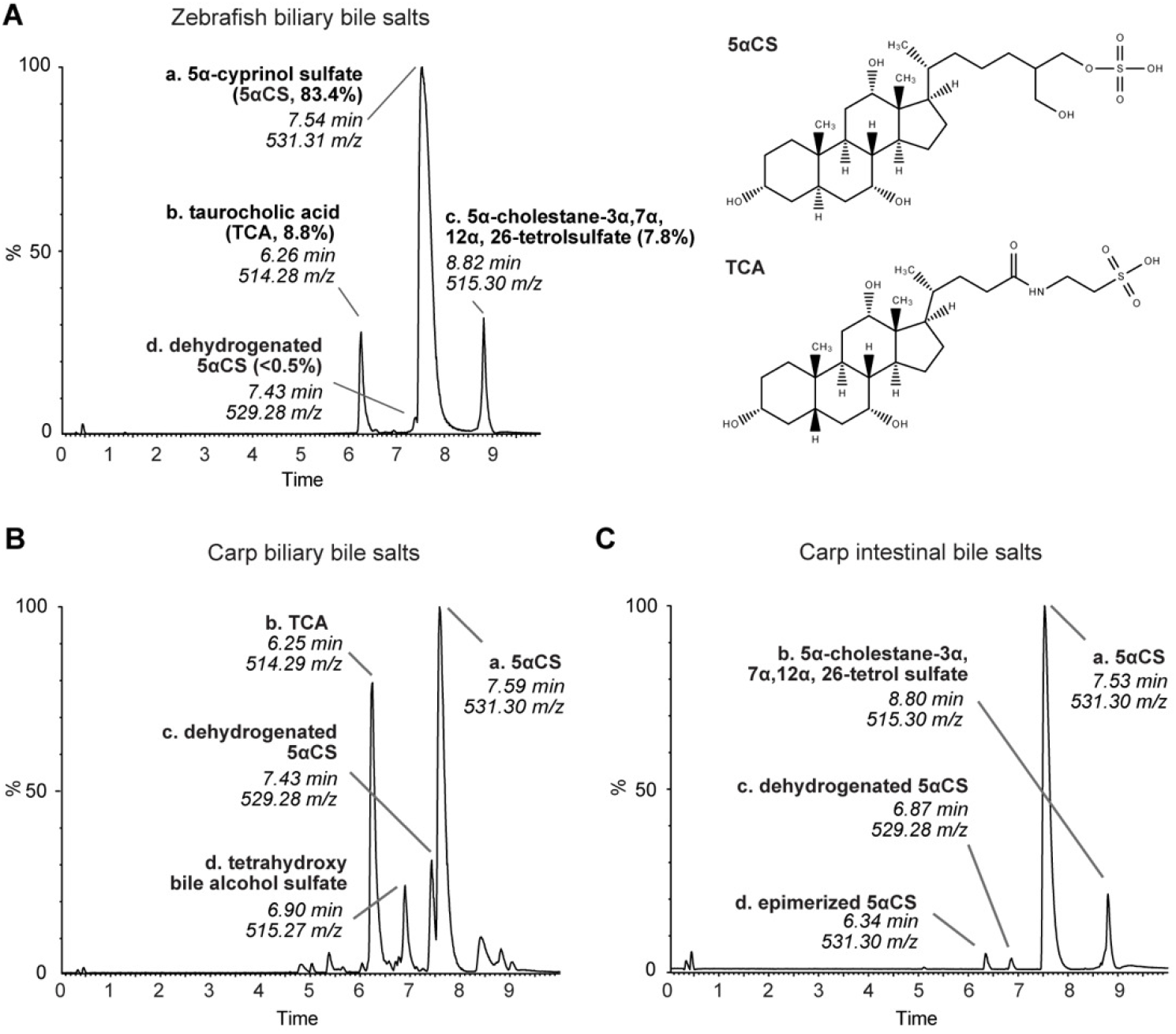
Zebrafish and carp bile salts consist of both C_24_ bile alcohols and C_27_ bile acids. **A.** Mass spectrometry chromatograms of the bile salts extracted from adult zebrafish gallbladders. The identity and proportion of major peaks are listed. The height of each peak is shown as the relative percentage that was normalized to the highest peak of the metabolites detected. Structures of 5αCS and TCA are shown. **B** and **C.** Mass spectrometry chromatograms of the bile salts extracted from adult carp gallbladders (B) or adult carp intestinal contents (C). The identities of major peaks are listed accordingly.

The predominant zebrafish bile salt, the C_27_ bile alcohol 5αCS, differs drastically from the common mammalian bile salts, the C_24_ bile acids, in both the stereostructure and the number of carbon atoms. Therefore, we asked whether such distinct bile salt composition results in differential regulation of Fxr signaling between fishes and mammals. We thus modulated the zebrafish bile salt levels by disrupting the hepatic synthesis or ileal uptake of bile salts and monitored the impacts on Fxr activity using the *Tg(-1.7fabp6:GFP)* reporter. To reduce hepatic bile salt synthesis, we generated a new *cyp7a1* mutant zebrafish (*cyp7a1^-16/-16^*, designated as *cyp7a1*^-/-^) which exhibited a significant reduction in the total bile salt levels as compared to its wt counterparts (Fig S1C-F, Supplementary Results). Using the reporter assay, we observed a over 50% decrease in GFP fluorescence in *cyp7a1* mutant zebrafish as compared to wt, suggesting that Fxr activity was reduced as a result of bile salt deficiency in zebrafish (Fig 1D). To reduce bile salt uptake in the ileum, we utilized *slc10a2* mutant (*slc10a2^sa2486/sa2486^*, designated as *slc10a2^-/-^*) zebrafish (Fig S1G) (*25*), which also showed significantly decreased ileal GFP fluorescence, consistent with compromised Fxr activity due to insufficient bile salt uptake (Fig 1E). Together, these results suggest that despite the compositional differences in bile salts between zebrafish and mammals, bile salts still activate Fxr and the downstream signaling in the ileal epithelium of zebrafish.

### Fish microbiota modulate bile salt diversity *in vivo* and *in vitro*

Primary bile salts can be modified by intestinal microbiota into various unconjugated or secondary bile salts and then cycled back to the liver through enterohepatic circulation (*5*). Our findings on zebrafish biliary bile salt diversity demonstrated the presence of a dehydrogenated 5αCS (Fig 2A). However, it is not clear if this modified 5αCS is an intermediate derived from *de novo* 5αCS biosynthesis or a recycled bile salt that has been modified by gut microbiota. Further, biliary bile salts do not accurately reflect the full spectrum of microbial modification that occur in the intestine. We therefore examined bile salt diversity in the intestinal contents of adult zebrafish, aiming to determine if microbial modifications of bile salts occur. 5αCS and TCA were present in zebrafish intestinal contents (Fig S2B); however, the low biomass of the zebrafish luminal contents limited our ability to accurately detect and/or quantify these bile salts and their derivatives. Thus, we turned to a larger cyprinid fish species closely related to zebrafish, the Asian grass carp (*Ctenopharyngodon idella*), and compared the bile salt diversity between the carp biliary bile and gut contents to determine if bile salts are modified by carp gut microbiota (Fig 2B, 2C). Carp possess a similar biliary bile salt profile to zebrafish, as all major peaks found in zebrafish were also present in carp (Fig 2A, 2B), except that it produces a different tetrahydroxy bile alcohol sulfate (Fig 2B, peak d), likely a 5β-isomer of the cholestane- 3α,7α, 12α, 26-tetrolsulfate (*1, 26*). Notably, in the bile salts isolated from carp intestinal contents, we observed a new peak sharing the same mass ratio but a different retention time with 5αCS, indicative of an epimerized 5αCS. This suggests that carp microbiota can oxidize and epimerize an α- to a β-hydroxyl group of the primary bile alcohol 5αCS (Fig 2C). To our knowledge, this is the first evidence demonstrating the ability of microbes to metabolize bile alcohols in vertebrates.

To test if similar and/or additional microbial modifications might be present in the zebrafish gut, we developed an *in vitro* bile salt modification assay using LC/MS (Fig 3A). Complex microbiota or individual microbes isolated from the zebrafish intestine were first enriched under aerobic or anaerobic conditions and then incubated with the bile salts of interest. This assay system was validated through successful detection of common modifications of primary bile salts upon treating with microbes known to perform these modifications (Fig S3A-B). We then used this system to test if zebrafish microbiota modify 5αCS and TCA, the primary bile alcohol and acid in zebrafish. For 5αCS cultured with zebrafish microbiota, two newly emerged peaks, representing the microbial metabolites of 5αCS, were detected in both aerobic and anaerobic conditions. One peak showed a mass ion of 529.3 m/z with an elution time of 7.4 min, suggesting a keto-5αCS variant (Fig 3B). The loss of two mass units observed in this product is consistent with bacterial hydroxysteroid dehydrogenase activity found in human gut microbiota (*27, 28*). The other peak, with a mass ion of 531.3 m/z and elution time of 6.4 min, corresponded to an epimerized 5αCS, a downstream product of the keto-5αCS variant, therefore further confirming the presence of hydroxysteroid dehydrogenases in zebrafish microbiota (Fig 3B). Interestingly, both peaks were also present in carp intestinal content (Fig 2C), suggesting that microbiota-mediated 5αCS dehydrogenation and epimerization are conserved in these cyprinid fishes.

**Figure 3.**
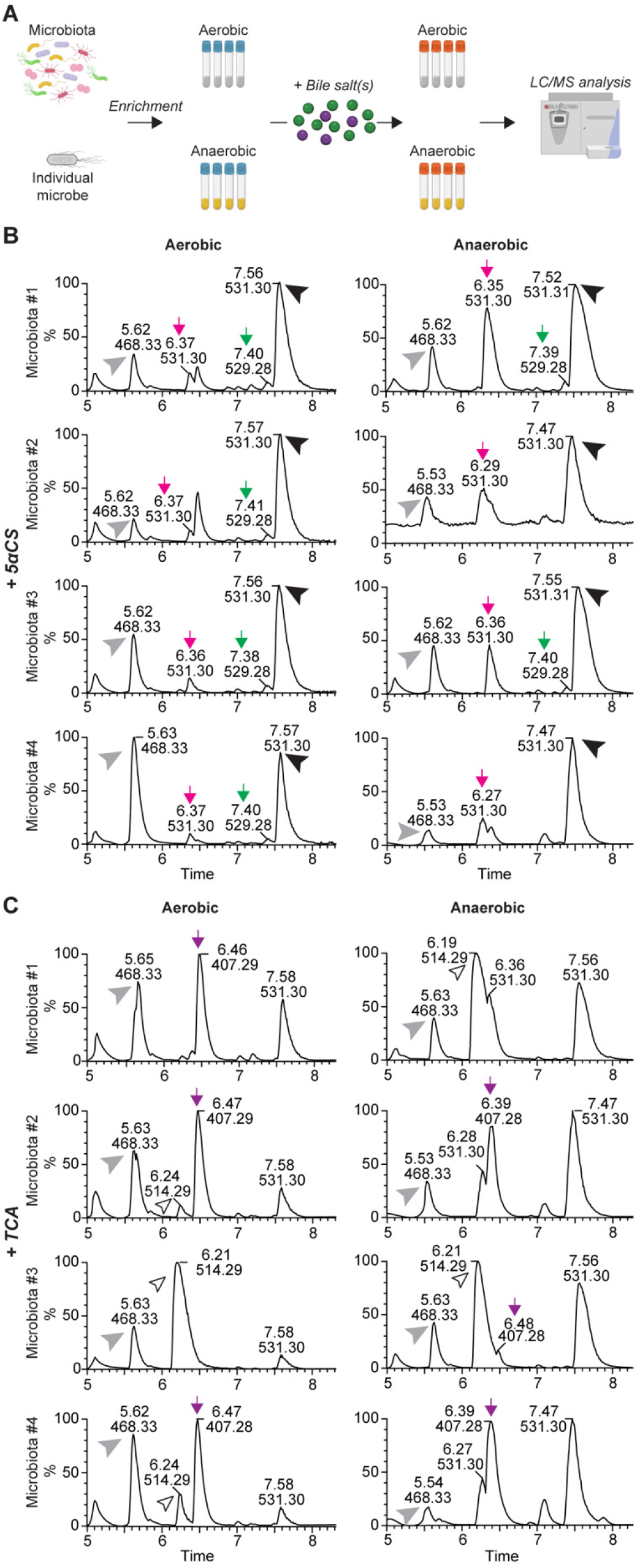
Microbiota modulate bile salt diversity in zebrafish. **A.** Schematic diagram of the *in vitro* bile salt metabolism assay. Complex microbiota or individual bacterial strains were enriched in modified TSB media under both aerobic and anaerobic conditions, and then incubated with 50 µ M of bile salts of interest, after which the total bile salts were extracted from individual bacterial cultures and subjected to LC/MS to identify modification of the supplemented bile salts. **B** and **C.** LC/MS chromatographs of bile salt metabolites extracted from enriched zebrafish microbiota cultures supplemented with 50 μM 5αCS (B) or TCA (C) under aerobic (left) or anaerobic (right) conditions. The arrowheads indicate the supplemented bile salts and internal control: black arrowhead: 5αCS; white arrowhead: TCA; grey arrowhead: internal standard D4-GCA. The arrows indicate bile salt metabolites resulted from microbial modification of the supplemented bile salts: green arrow: dehydrogenated 5αCS; magenta arrow: epimerized 5αCS; purple arrow: CA. The height of each peak is shown as the relative percentage that was normalized to the highest peak of the metabolites detected. Shown are representative chromatographs of zebrafish microbiota from four independent experiments.

Cultures incubated with TCA resulted in a new peak corresponding to cholic acid (CA), the deconjugated product of TCA, in several zebrafish microbiota, though the extent of deconjugation varied (Fig 3C). For example, aerobic microbiota #1 exhibited a complete deconjugation of TCA to CA whereas aerobic microbiota #3 showed no sign of deconjugation. This likely indicates the variable distribution of microbes containing bile salt hydrolase (BSH), the enzyme catalyzing the deconjugation of TCA, among zebrafish. Interestingly, the presence or absence of BSH activity in a given zebrafish microbial community does not always match between aerobic and anaerobic conditions. For instance, zebrafish microbiota #3 deconjugated TCA only under anaerobic conditions, whereas microbiota #1 catalyzed deconjugation only under aerobic conditions. This suggests that the bacteria responsible for deconjugation are likely different among distinct zebrafish microbiota and that more than one deconjugating bacterium is present in zebrafish. No other transformations of 5αCS or CA were detected in cultures. Collectively, our results indicate that microbial modification of bile salts is a conserved feature between zebrafish and mammals.

Having shown that zebrafish microbiota modify both 5αCS and TCA, we sought to determine the bacterial specificity of bile salt modification in zebrafish. We screened a panel of zebrafish gut isolates representing several major bacterial taxa in the zebrafish gut towards 5αCS and TCA (Fig 4A, S3C). None of the tested strains modified 5αCS. Yet, we identified one Gammaproteobacteria strain, *Acinetobacter sp.* ZOR0008, capable of deconjugating TCA (Fig 4A). After overnight incubation with *Acinetobacter* sp., we observed complete conversion of 25 μM TCA to CA, suggesting robust BSH activity (Fig 4A). To our knowledge, this the first zebrafish gut bacterium confirmed to have bile salt metabolizing activity.

**Figure 4.**
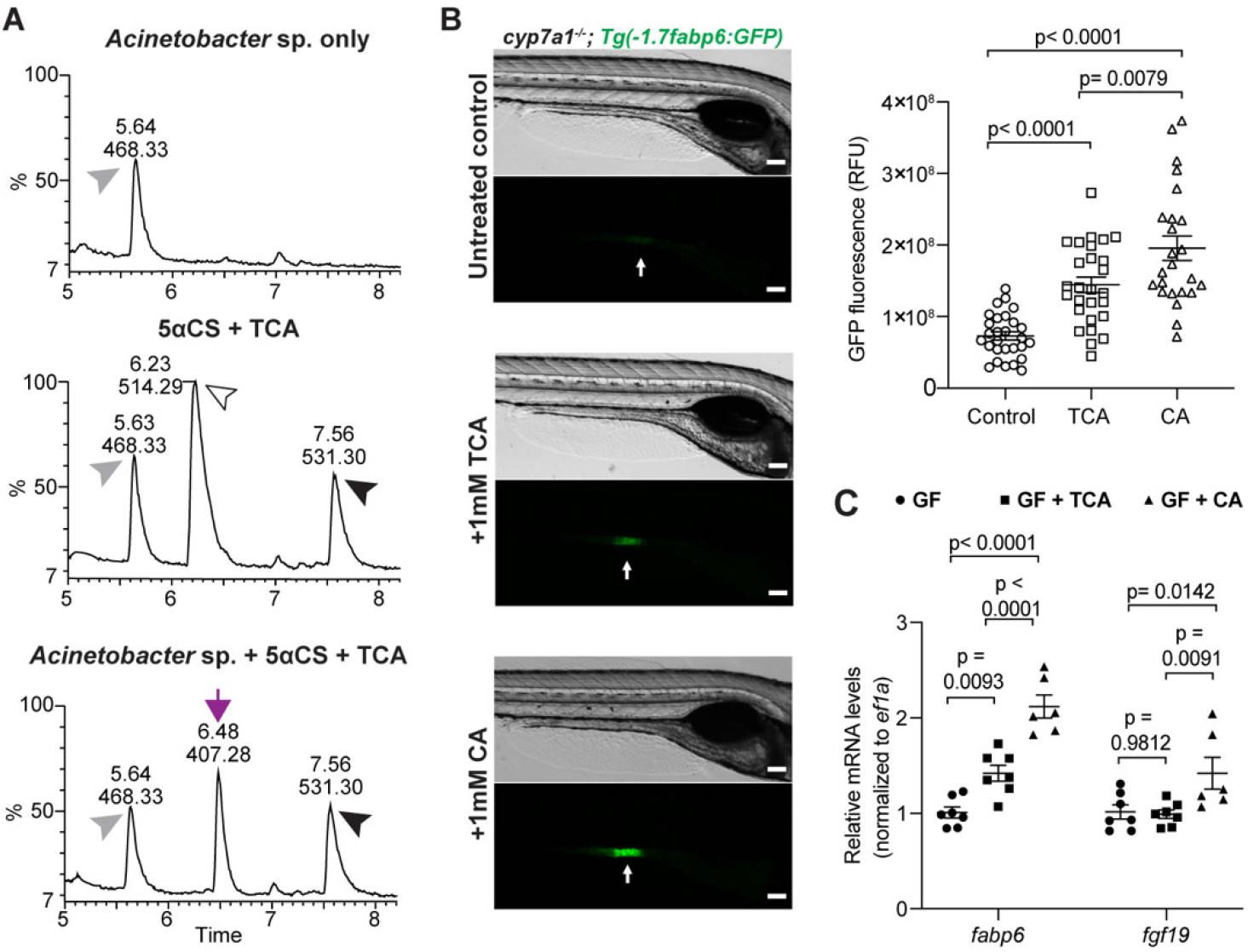
A zebrafish gut bacterium modifies primary bile acid producing a metabolite that augments Fxr signaling in zebrafish. **A.** LC/MS chromatographs of medium only control (upper), bile salts only control (middle), and *Acinetobacter* sp. ZOR0008 culture supplemented with bile salts (bottom) under aerobic conditions. The arrowheads indicate the supplemented bile salts and internal control: black arrowhead: 5αCS; white arrowhead: TCA; grey arrowhead: internal standard D4-GCA. The purple arrow indicates the liberated CA resulted from bacterial deconjugation of TCA. Results from other bacterial strains are shown in Fig S3C. **B.** Imaging and quantification of GFP fluorescence of the ileal region of 7 dpf *Tg(-1.7fabp6:GFP) cyp7a1*^-/-^ larvae that were untreated or treated with either 1 mM TCA or CA for 4 days. The zebrafish ileal region is indicated by arrows. Scale bar = 100 μm. **C.** qRT-PCR comparing the expression of Fxr target genes in 7 dpf wt germ free larvae that were untreated or treated with either 1mM TCA or CA for 4 days. The results are represented as relative expression levels normalized to the housekeeping gene *ef1a* (Mean±SEM). Statistical significance was calculated by one-way (B) and two-way (C) ANOVA with Turkey’s multiple comparisons test. Shown are representative data from at least three independent experiments.

### Microbial modifications of bile salt in zebrafish modulate Fxr activity

In mammals, microbial modification of bile salts can alter the signaling property of bile salts and modulate host physiology. Given that the primary bile acid TCA can be metabolized into CA by gut microbes in zebrafish, we investigated the potential impact of that modification on Fxr signaling *in vivo* by monitoring *Tg(-1.7fabp6:GFP)* zebrafish treated with exogenous TCA or CA. To reduce the influences of Fxr activity caused by endogenous bile salts, we performed the reporter assay in the *cyp7a1^-/-^* background. Physiological concentrations of TCA or CA were supplemented to larval zebrafish and GFP fluorescence was monitored after 4 days (*23*). Zebrafish larvae treated with CA exhibited increased GFP fluorescence as compared to those with TCA, and both showed higher fluorescence than the non-treated controls (Fig 4B). This suggests that both TCA and CA activate Fxr and that CA is more potent than TCA, consistent with observations in mammals (*29*). We further validated these findings under a more stringent setting using wt germ-free zebrafish, which permit competition between endogenous versus exogenous bile salts and eliminate potential influences of microbiota on metabolizing the supplemented bile salts. Quantitative RT-PCR (qRT-PCR) results suggested that CA treatment increased expression of Fxr targets, such as *fabp6* and *fgf19*, as compared to TCA, confirming that CA displays higher potency than TCA in activating Fxr (Fig 4C). Together, our observations suggest that zebrafish gut microbiota have the potential to regulate Fxr-mediated signaling through modification of primary bile salts.

### Fxr regulates diverse cell types identified in zebrafish intestine by single-cell RNA-seq

Having established that bile salts and gut microbes interactively regulate Fxr activity, we next sought to discern how Fxr in turn contributes to intestinal functions. Gross intestinal morphology appeared normal in zebrafish and mice lacking Fxr function (Fig 1B) (*30*), but strong attenuation of the *fabp6* reporter in *fxr* mutant zebrafish (Fig 1B) suggested potential effects of *fxr* mutation on functional specification of intestinal epithelial cells (IECs). To test this possibility, we performed single-cell RNA sequencing (scRNA-seq) on 12,543 IECs sorted from 6 dpf *fxr*^+/+^ or *fxr*^-/-^ zebrafish larvae on a *TgBAC(cldn15la-GFP)* transgenic background that expresses GFP in all IECs (*31*). After quality control, 4,710 cells from *fxr*^+/+^ and 5,208 cells from *fxr^-/-^* samples were used for downstream analyses (Fig S4A-E). Twenty-seven distinct clusters were generated by unsupervised clustering of these cells using the Seurat R package as described previously (Fig 5A, S4A-E) (*32*). The cell types represented by these clusters were inferred through integrative analysis of published expression data of previously identified gene markers, novel markers of each cluster identified in this study by differential gene expression, and functional predictions from the gene expression data generated in this study (Datasets S1-3, and Supplementary Results). The resulting annotation revealed a range of IEC types including absorptive enterocytes, goblet cells (including those that resemble mammalian tuft cells and microfold cells), enteroendocrine cells, secretory precursors, ionocytes (including those that resemble mammalian BEST4/OTOP2 cells) (*33*), and foregut epithelial cells, as well as low levels of several other apparent contaminating cell types (e.g., exocrine pancreas cells, epidermis cells, mesenchymal cells, leukocytes, red blood cells) (Fig 5A, Table S1, and Supplementary Results). These results combined with our extended annotation of this scRNA-seq dataset provided in Supplementary Materials provide a useful new resource for zebrafish intestinal biology.

**Figure 5.**
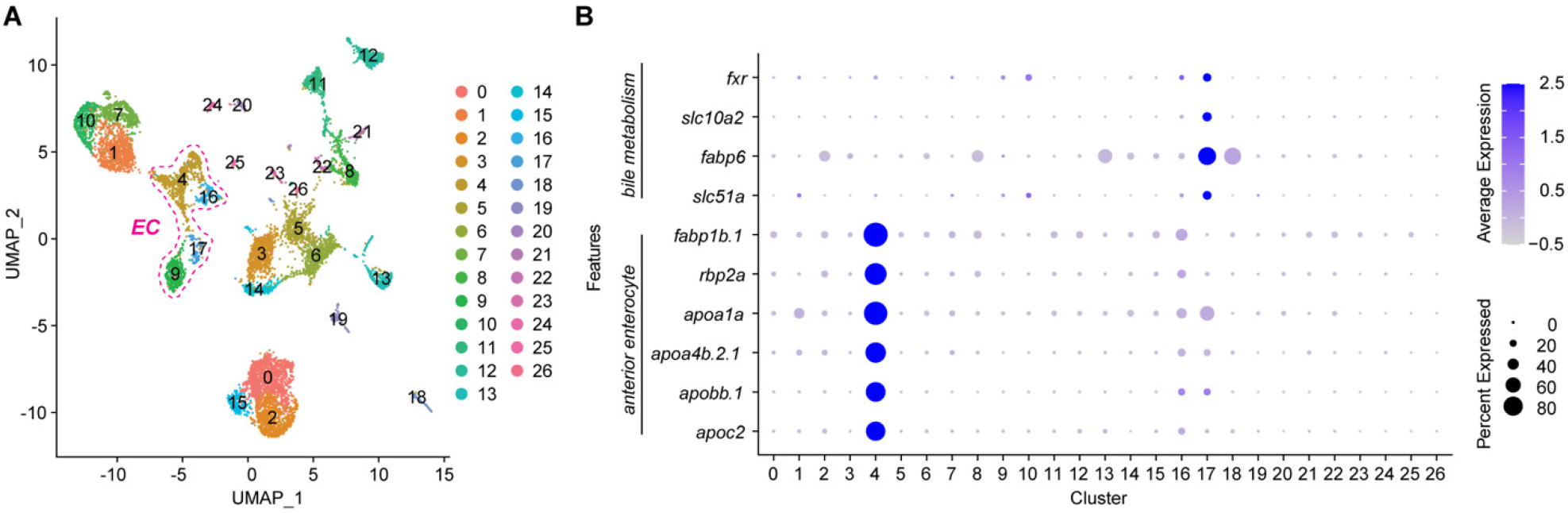
ScRNA-seq reveals extensive cellular diversity in the larval zebrafish intestinal epithelium. **A.** UMAP visualization of cell-type clusters using combined *fxr*^+/+^ and *fxr*^-/-^ single-cell transcriptomes (4,710 cells from *fxr*^+/+^ and 5,208 cells from *fxr*^-/-^). Clusters were hierarchically ordered in PCA spaces and were numbered accordingly. Detailed annotations of each cluster are shown in Table S1 and Supplementary Results. Clusters 4, 16, 17, and 9 are absorptive enterocytes (EC). **B.** Dot plots showing relative expression of bile salt metabolism genes and anterior enterocyte markers in *fxr*^+/+^ cells throughout all clusters.

We next leveraged our scRNA-seq data to test the requirement for *fxr* in different IEC types. Supporting the notion that Fxr regulates diverse aspects of intestinal physiology, we found that nearly one-third of all clusters exhibited over 50% change in cell abundance with an average of ∼500 genes displaying over 1.5-fold changes in expression in response to Fxr mutation (Fig S5A, B, Dataset S4). To further evaluate conservation of Fxr-mediated gene expression between zebrafish and mammals, we compared these results to an existing dataset of 489 mouse genes differentially regulated in the ileum or colon in response to intestinal Fxr agonism (*34*). We identified 583 zebrafish genes that were determined by BioMart to be homologous to those 489 mouse genes and also detected in our zebrafish dataset. Of the 583 zebrafish genes, 213 of them were differentially expressed in response to *fxr* mutation in at least one cluster (Dataset S5). In those instances where one of those 213 genes was differentially expressed in a cluster, the directionality of change due to Fxr function was consist with mouse ileum in 59.5% (72/121) of cases and with mouse colon in 49.7% (251/499) of cases. Though it remains unknown if these gene expression changes are due to direct or indirect effects of Fxr activity, these results do suggest substantial differences between the gene regulons influenced by Fxr activity in the zebrafish and mouse intestines. This further underscores the importance of using multiple animal models to gain complementary insights into bile salt-Fxr signaling pathways. Although we already showed that loss of *fxr* function in zebrafish results in reduction of several conserved Fxr target genes (Fig 1C), this comparative functional genomic analysis identified potential additional targets of Fxr regulation that are conserved between zebrafish and mice such as *Pck1* (*35*), *Akr1b7* (*36*), and *Apoa1* (*37*) (Dataset S5). Collectively, our scRNA-seq results unveil extensive cellular diversity in the larval zebrafish intestine and highlight the broad impacts of Fxr on cell abundance and gene expression in diverse cell types.

### Fxr regulates functional specialization of ileal epithelial cells

Given the striking attenuation of the ileal *fabp6* reporter in *fxr* mutant zebrafish (Fig 1B), we next examined how Fxr impacts zebrafish ileal epithelial cells in our scRNA-seq analysis. We discerned cluster 17 as enriched for zebrafish ileal epithelial cells based on the abundant expression of bile transporters *fabp6*, *slc10a2*, and *slc51a* (Fig 5B). This cluster exhibited a higher level of *fxr* expression as compared to all other clusters, consistent with the notion that *fxr* displays spatially patterned expression along the intestine with highest levels in the ileal epithelium (*38, 39*). We also observed heterogeneity in the expression of these bile transporter genes in cluster 17, raising the possibility that multiple sub-cell types are present in this cluster (Fig S6A). For clarity, we operationally defined cells located in cluster 17 as “ileal epithelial cells” and the subset that expresses one or more bile transporters as “ileocytes”.

Mutation of *fxr* drastically impacted gene expression in cluster 17 cells (Fig S5B, Dataset S4). As expected, many downregulated genes in cluster 17 *fxr* mutant cells were related to bile salt metabolism, such as *fabp6*, *slc10a2*, and *slc51a*, consistent with the strong reduction in the *fabp6* reporter activity upon Fxr mutation (Fig 6A, 1B). On the other hand, among the upregulated cluster 17-enriched markers, lysosome process (dre: 04142) was the most enriched pathway in *fxr* mutant cells (Fig 6A, S6B). Lysosome mediated degradation is a hallmark function of a specific type of vacuolated enterocytes named lysosome-rich enterocytes (LRE) (*40*). LREs are found in the ileum of fishes and suckling mammals and are known to internalize dietary macronutrients for intracellular digestion (*41, 42*). Therefore, our data confirm that LREs compose a key type of ileal epithelial cell, and further suggest that Fxr normally represses LRE gene expression. Indeed, we observed increased expression of many known LRE makers in *fxr* mutant cells in cluster 17. This includes multiple classes of digestive enzymes involved in macromolecule degradation and transporters responsible for dietary protein uptake in LREs (Fig 6A) (*40*). To confirm these results, we used qRT-PCR to examine expression of several LRE markers including *amn*, which encodes Amnionless, the major component of the multi-ligand endocytic machinery in LREs, and *ctsbb*, which encodes Cathepsin B commonly found in lysosomes. Both genes exhibited higher expression in the zebrafish intestine upon *fxr* mutation, validating our scRNA-seq observations (Fig 6B). To identify potential transcriptional regulatory pathways involved in the induction of LRE genes upon *fxr* mutation, we searched for transcription factor binding sites (TFBS) that are over-represented within accessible chromatin regions (*17*) near genes upregulated in *fxr* mutant cells in cluster 17. The top 3 enriched TFBS were ZBTB33, Atf2, and TATA-box, raising the possibility that Fxr may interact with TFs that bind at these TFBSs to regulate LRE functions such as lysosomal mediated degradation (Fig S6C). Collectively, these results establish that Fxr promotes expression of bile absorption genes and represses expression of lysosomal degradation genes in ileal epithelial cells in zebrafish.

**Figure 6.**
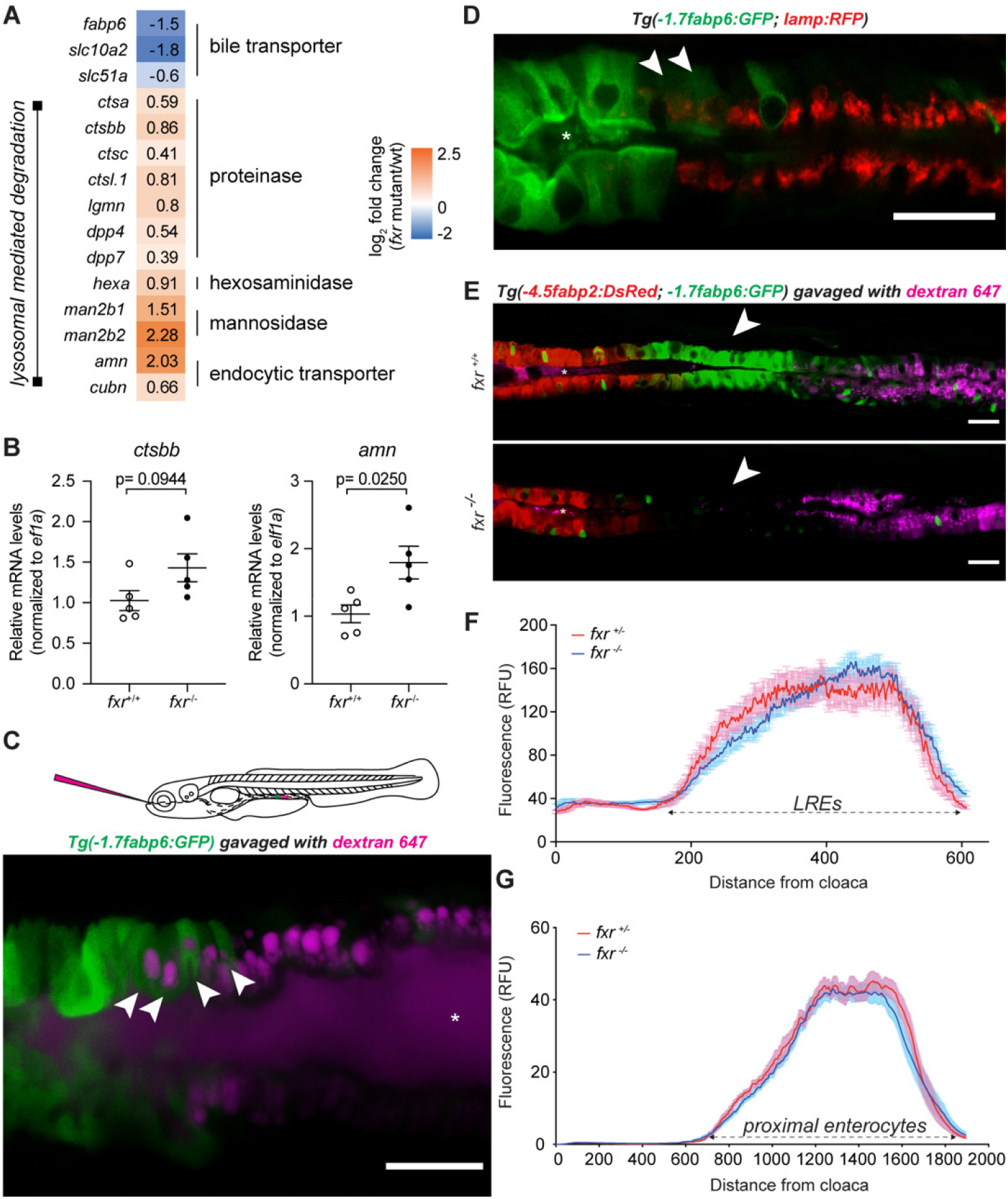
Fxr determines specific cell functions of ileal epithelial cells but is not required for establishment of regional boundaries. **A.** Differential expression of genes related to bile metabolism and lysosomal functions in *fxr*^-/-^ cells as compared to *fxr*^+/+^ cells in cluster 17. **B.** qRT-PCR comparing expression of LRE markers in dissected intestines of gender and size-matched adult *fxr*^+/+^ or *fxr*^-/-^ zebrafish. The results are represented as relative expression levels normalized to the housekeeping gene *ef1a* (Mean±SEM). Statistical significance was calculated by unpaired t-test. Shown are representative data from 2 independent experiments. **C.** Confocal single-plane image of the intestinal epithelium in 7 dpf live larvae expressing the *Tg(-1.7fabp6:GFP)* transgene (green) following gavage with Alexa Fluor 647 Dextran (magenta). White arrowheads mark GFP- expressing cells that uptake dextran. The schematic diagram depicting gavage approach for labeling lysosome rich enterocytes (LREs) in the larval zebrafish intestine using fluorescent luminal cargo is shown in the top panel. Image is representative of 6 larvae examined. **D.** Confocal three-dimensional projections of the intestinal epithelium of 7 dpf larvae expressing the *Tg(-1.7fabp6:GFP)* (green) and the *TgBAC(lamp2:RFP)* (red) transgenes. White arrowheads mark cells showing both GFP and RFP expression. Image is representative of 5 larvae examined. **E.** Confocal single-plane image of the intestinal epithelium in 7 dpf *fxr*^+/+^ (top) and *fxr*^-/-^ (bottom) larvae expressing the *Tg(-4.5fabp2:DsRed*) (red) and *Tg(-1.7fabp6:GFP)* (green) transgenes following gavage with Alexa Fluor 647 Dextran (magenta). White arrowheads mark the ileocyte region, which persists in *fxr*^-/-^ larvae despite loss of GFP expression. Images are representative of 5 *fxr*^+/+^ and 5 *fxr*^-/-^ larvae examined. Asteroids mark zebrafish lumen in (C, D, E). Scale bar = 25 μm. **F.** Uptake profiles along LRE region following gavage with 1.25 mg/mL Alexa Fluor 647 Dextran in 7 dpf *fxr*^+/-^ (n= 14) and *fxr*^-/-^ (n= 22) larvae. G. DsRed fluorescence along the intestine of 7dpf *fxr*^+/-^ (n= 18) and *fxr*^-/-^ (n= 24) *Tg(-4.5fabp2:DsRed)* larvae.

The altered gene expression seen in *fxr* mutant cells in cluster 17 could be explained by Fxr regulating the relative abundance of different ileal cell types such as ileocytes and LREs, or regulating expression of genes characteristic of those cell types in cluster 17. We therefore carried out subclustering of cluster 17, aiming to distinguish ileocytes from LREs and to delineate the heterogeneity within these ileal epithelial cells (Fig S6D). To our surprise, we could not cleanly separate these two subtypes as many cluster 17 cells expressed both bile transporter genes and LRE makers (Fig S6D-F). This indicates an overlap between the bile salt absorption and lysosomal degradation programs in some cluster 17 ileal epithelial cells, and is in agreement with previous bulk RNA-seq studies showing that LREs can also express ileocyte markers such as *fabp6* and *slc10a2* (*40*). To test this overlap *in vivo*, we took advantage of the high endocytotic property of LREs and labeled them in *Tg(-1.7fabp6:GFP)* zebrafish by gavaging with fluorescent dextran which is internalized by LREs (Fig 6C) (*40*). Indeed, some ileal epithelial cells were labeled by both GFP and dextran, whereas cells anterior to this region were only GFP+ and cells posterior to this region were only dextran+. This was confirmed with a second LRE reporter *TgBAC(lamp2:RFP)* (Fig 6D), further establishing the partial overlap of these two functionally distinct transcriptional programs. Collectively, these findings demonstrate that cluster 17 represent cells located in the zebrafish ileum that include at least three subtypes that we operationally define as (1) ileocytes, which express bile metabolism genes and are responsible for bile salt absorption; (2) LREs, which express lysosomal enzymes and are responsible for macromolecule degradation; and (3) bi-functional cells exhibiting both of those programs.

We next sought to determine if Fxr regulates the abundance or location of these ileal cell types. In contrast to the striking changes in the gene expression of cluster 17 cells upon *fxr* mutation (Fig 6A, S5B), the relative abundance of this cluster remained similar between *fxr* mutant and wt (Fig S5A), suggesting that Fxr deficiency does not prevent the establishment of a zebrafish ileum. To validate this observation *in vivo*, we measured the length and the spatial location of the ileal epithelium, including ileocytes and LREs, in *fxr* wt or mutant zebrafish. The LRE region was evaluated by gavaging dextran into *fxr* wt and mutant zebrafish followed with *in vivo* imaging. No significant difference was observed in the length of the dextran positive region or the intensity of the absorbed dextran after gavaging, suggesting that the abundance of LREs remain unchanged in the absence of Fxr (Fig 6E, F). To assess ileocyte abundance and positioning, we gavaged dextran into the double transgenic reporter *Tg(-4.5fabp2:DsRed, - 1.7fabp6:GFP)* in either *fxr* wt or mutant background (*43*). The proximal intestinal region, labeled by DsRed, demarcates the anterior boundary of the ileocytes, while the LRE region, labeled by dextran, demarcates the posterior boundary. The anterior boundary remained intact in the *fxr* mutant zebrafish, as the DsRed region did not expand or contract (Fig 6E, G). We did observe a nearly complete loss of GFP fluorescence in the *fxr* mutant animals, consistent with the findings that *fabp6* is under strong regulation by Fxr (Fig 6E, 1A). However, this non-fluorescent region, flanked by the anterior enterocytes and LREs in the *fxr* mutant, shares similar length and spatial position as the GFP positive region in the *fxr* wt zebrafish (Fig 6E-G). These findings are in agreement with our scRNA-seq data and confirm that Fxr impacts the gene expression program of the ileal epithelial cells without overtly affecting the proportion of those cells, nor the segmental boundaries that organize that region of the intestine. Together, our data suggest that Fxr is not required for developmental organization of the ileal region, instead it is involved in distinct physiological aspects of the cell types in this region.

### Fxr promotes differentiation of anterior absorptive enterocytes

Since *fxr* is expressed along the length of the intestine in zebrafish and mammals (Fig 5B) (*38*), we next examined how Fxr contributes to the functions of absorptive enterocytes other than the ileal epithelial cells. We focused on cluster 4, which represents mature enterocytes in the anterior intestine based on their expression of known jejunal markers such as *fabp1b.1* and *rbp2a*, as well as genes involved in lipid metabolism, a hallmark function of mammalian jejunum (Fig 5B, Table S1, Datasets S2-3) (*44*). Our scRNA-seq data suggested increased cell abundance of cluster 4 in response to Fxr mutation (Fig S5A). To validate this *in vivo*, we performed fluorescence-activated cell sorting of anterior enterocytes collected from double transgenic *fxr* wt or mutant fish harboring the anterior enterocyte reporter *Tg(-4.5fabp2:DsRed)* and the pan-IEC reporter *TgBAC(cldn15la-GFP)*. Consistent with our scRNA-seq data, we observed a significant increase in the relative abundance of anterior enterocytes in the *fxr* mutant compared to wt (Fig 7A), confirming the role of Fxr in regulating the abundance of these cells in zebrafish. Fxr mutation also led to altered expression of over 250 genes in cluster 4 cells (Fig S5B, Dateset S4). Interestingly, the majority (∼86%) of these differentially expressed genes were downregulated. Functional categorization analysis revealed that these downregulated genes in *fxr* mutant cells in cluster 4 were enriched for pathways involved in energy metabolism of diverse substrates (Fig S7). This includes aspects of lipid metabolism such as lipid biosynthesis (GO term: sterol biosynthetic process), trafficking (GO term: plasma lipoprotein particle assembly), and regulation (dre: PPAR signaling pathway), amino acid metabolism (GO terms: peptide metabolic process; cellular modified amino acid metabolic process; creatine metabolism), and xenobiotic metabolism (GO terms: drug metabolic process; response to xenobiotic stimulus). Since these pathways represent key functions of differentiated anterior enterocytes, we speculated these gene expression differences in *fxr* mutant cells in cluster 4 may be due to reduced differentiation of these enterocytes. We therefore compared the zebrafish genes differentially regulated in *fxr* mutant cells in cluster 4 against defined sets of signature genes for intestinal stem cells (ISCs) and differentiated enterocytes in the small intestinal epithelium of adult mice (Fig 7B) (*45*). This revealed an overlap of 102 one-to-one gene orthologs between the downregulated genes of the *fxr* mutant cells in cluster 4 in this study and the genes preferentially expressed in either ISCs or differentiated enterocytes in mice. Approximately two-thirds of these genes (64 out of 102) are preferentially expressed in differentiated enterocytes, suggesting that Fxr inactivation in cluster 4 preferentially attenuates enterocyte differentiation programs. In support, we observed that the most enriched TFBS within accessible chromatin near the genes downregulated in *fxr* mutant anterior enterocytes is Hnf4α, a TF known to promote enterocyte differentiation (Fig 7C) (*46, 47*). Collectively, these results reveal a novel role of Fxr in promoting differentiation programs of anterior enterocytes.

**Figure 7.**
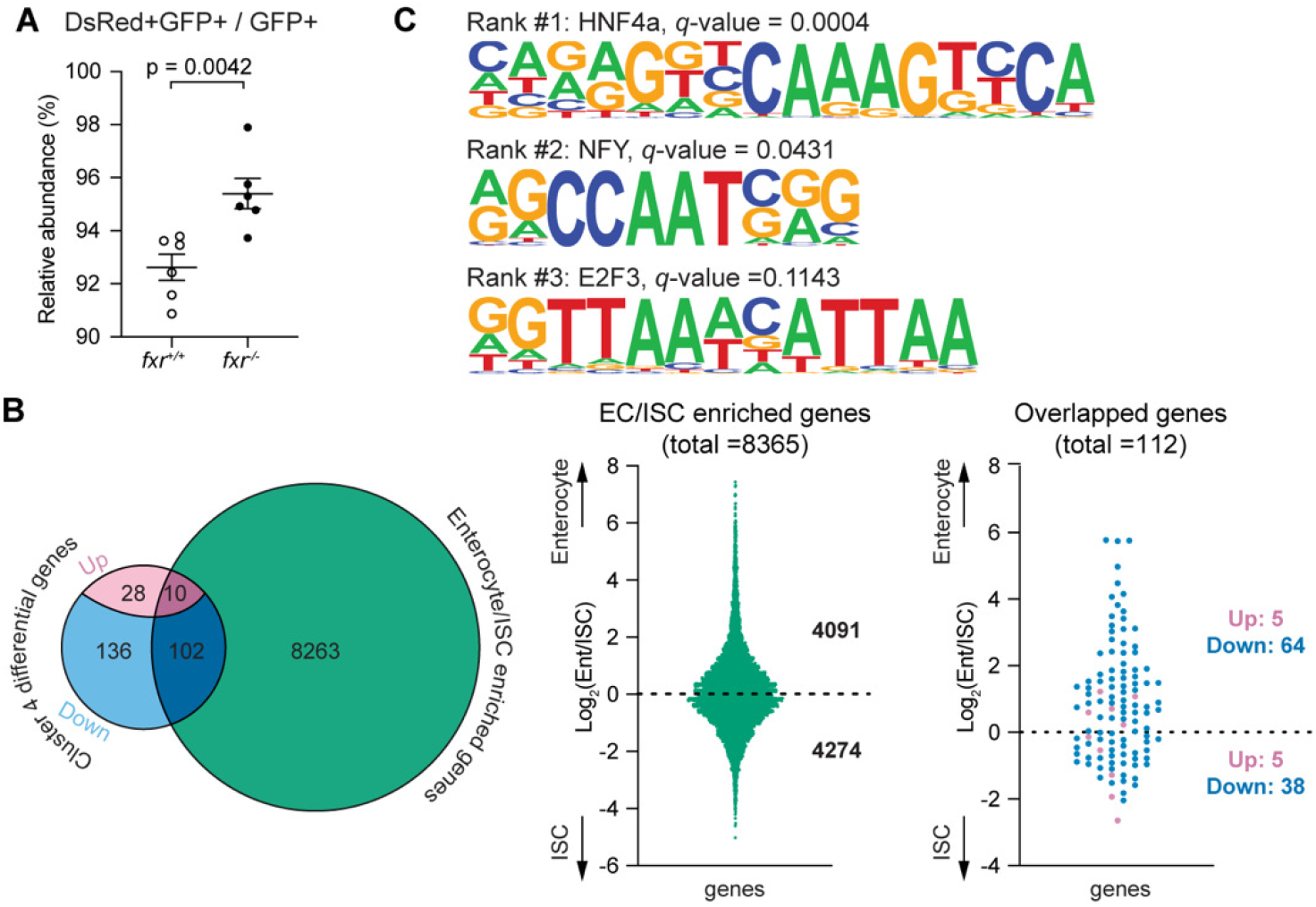
Fxr regulates differentiation and functions of anterior absorptive enterocytes in zebrafish. **A.** FACS analysis comparing the relative abundance of anterior enterocytes in 7 dpf *fxr*^+/+^ and *fxr*^-/-^ larvae expressing the *Tg(-4.5fabp2:DsRed)* and *TgBAC(cldn15la-GFP)* transgenes. The relative abundance was calculated by dividing the cell counts of DsRed+GFP+ double positive cells by the cell counts of GFP+ single positive cells (Mean±SEM). Statistical significance was calculated by unpaired t-test. Shown are representative data from two independent experiments. **B.** Comparisons between the genes that were differentially expressed in *fxr*^-/-^ cells relative to *fxr*^+/+^ cells in cluster 4 and the mouse genes enriched in jejunal enterocytes/intestinal stem cells (ISC) (*45*). Left: Venn diagram showing the number of overlapped genes between the two gene sets, of which the differentially expressed genes in *fxr*^-/-^ cells were further classified based on the changes in their expression (up- or down-regulated) upon *fxr* mutation. Middle: The distributions of enterocyte- and ISC-enriched genes from the enterocyte/ISC dataset used in the current comparison. Right: The distribution of overlapped genes resulting from the comparison. **C.** The top 3 HOMER-identified motifs enriched within accessible chromatin regions near genes that were down regulated in the *fxr*^-/-^ cells relative to *fxr*^+/+^ cells in cluster 4. Shown are the position weight matrices (PWMs) of the enriched nucleotide sequences. The TF family that most closely matches the motif is indicated above the PWM.

## Discussion

The ability to synthesize bile salts and the bile salt-regulated transcription factor Fxr are common features of all vertebrate classes, yet our knowledge of bile salt metabolism and bile salt-Fxr signaling is largely derived from mammals. Here, we characterize the bile salt-Fxr signaling axis in zebrafish by determining the bile salt composition and the key genetic components of Fxr signaling pathways. Further, we elucidate the microbiota-bile salt-Fxr relationships in zebrafish and highlight the importance of these interactions as they have been conserved over 420 million years since the last shared common ancestor between mammals and fishes. Collectively, we establish zebrafish as a valuable non-mammalian vertebrate model to study the bile salt-Fxr signaling axis and host-microbe coevolution. Using this model, we uncover novel functions of Fxr in modulating transcriptional programs controlling regional metabolic activities in the zebrafish intestine, including its role in repressing genes important for LRE functions in the ileum and promoting genes involved in enterocyte differentiation in the anterior intestine.

Our data show that zebrafish bile salts are composed predominantly of the evolutionarily “ancestral” C_27_ bile alcohol 5αCS, with only a small proportion of the evolutionarily recent C_24_ bile acid TCA that are commonly found in mammals (Fig 2A). To our surprise, zebrafish 5αCS was not observed to undergo 7α-dehydroxylation, a common microbial modification of bile acids in mammals (Fig 3B, S3B-C) (*48*). Further, we did not observe 7α-dehydroxylation of CA by the zebrafish or carp microbiota, even though CA is a suitable substrate for this modification in the mammalian gut environment (Fig 2C, 3C). Interestingly, although mammalian gut microbes have evolved numerous sulfatases that recognize and hydrolyze bile acid-sulfates (*49, 50*), we did not detect sulfatase activity towards 5αCS in zebrafish microbiota (Fig 3B). Oxidation and epimerization of bile acids by microbial hydroxysteroid dehydrogenase enzymes is well documented in mammalian gut microbiota (*27, 51, 52*) and is now confirmed to occur in zebrafish (Fig 3C). Deconjugation of TCA by bacterial BSHs was also observed in the present study (Fig 3C, 4A), a function widespread among mammalian microbial taxa (*53*), and important for regulation of lipid and cholesterol metabolism in diverse vertebrates (*54*). Together, these findings indicate that there may be top-down selection pressure in zebrafish to prevent evolution or acquisition of microbial enzymes that would recognize the side-chain sulfate and/or the 7α- hydroxyl group, therefore limiting secondary bile alcohol/acids production. Future study on how these primary and secondary bile salts contribute to digestive physiology and host-microbe interactions in different animals will shed light on understanding the evolutionary biology of vertebrate bile salts.

The binding pocket of FXR and the bile salt structures within a given vertebrate species are thought to have a co-evolutionary relationship (*7, 19*). For example, C_27_ bile alcohol 5αCS, the major bile salt species in zebrafish, specifically binds and activates zebrafish Fxr but not mammalian FXR (*7*). Here, we show that the C_24_ bile acids TCA, a minor zebrafish bile salt, along with its derivative CA, can both stimulate zebrafish Fxr activity *in vivo* (Fig 4B-C). This raises the possibility that zebrafish Fxr structure is able to bind both the ancestral bile alcohols and the modern bile acids, thereby representing an evolutionary transitional state. Additionally, we find that key aspects of the Fxr signaling pathways remain conserved between zebrafish and mammals including Fxr-mediated induction of *fabp6* and *fgf19* (Fig 1C). We further show that these zebrafish Fxr-dependent genes, like their mammalian homologs, are more potently induced by the microbially-derived deconjugated bile acid CA compared to its primary bile acid precursor TCA (Fig 4B, C) (*29*). Beyond these similarities, our analysis of intestinal genes regulated by Fxr function in zebrafish and mice also revealed significant differences. An important example is the directionality of Fxr regulation of *slc10a2*. Unlike in humans and mice where FXR represses *Slc10a2*, Fxr in zebrafish appears to induce *slc10a2*, as both the *fxr* and *cyp7a1* mutants displayed reduced *slc10a2* expression (Fig 1C, S1F). This suggests divergence of regulation of *slc10a2* by Fxr since the common ancestor of fish and mammals. In fact, Fxr-mediated regulation of *Slc10a2* homologs differs considerably even within mammals (*55*). For example, FXR negatively regulates intestinal *Slc10a2* in mice but not in rats (*56, 57*). Further, although *Slc10a2* is repressed by FXR in both mice and humans, the underlying mechanisms are different (*58*). Future structure-function analyses are warranted to dissect the regulatory mechanisms responsible for such differential control of *slc10a2* as well as the physiological consequences.

The zebrafish intestinal epithelial cell scRNA-seq dataset reported here provides a useful new resource for zebrafish intestinal biology. The perspectives afforded by this scRNA-seq dataset allowed us to evaluate distinct regulatory roles of Fxr across different intestinal cell types. For example, we found that ileal epithelium (identified as cluster 17 in this dataset) is composed of multiple cell subtypes including ileocytes, LREs, and bifunctional cells expressing both bile transporter genes and lysosomal degradation markers (Fig 6, S6E-F). Close relationships between ileocytes and LREs have also been defined in mammals, suggesting they are ancient cellular features of the vertebrate ileum. LREs develop in the mammalian ileum only during suckling stages before being replaced by ileocytes post-weaning (*40, 59*). Expression of genes involved in lysosomal degradation declines during this transition, whereas the expression of genes associated with bile salt absorption increases, suggesting an inverse correlation between these two functions (*40*). Our results provide potential mechanistic insight into the regulation of these two functions by demonstrating that Fxr promotes the expression of bile salt absorption genes and concomitantly reduces lysosomal degradation genes in the zebrafish ileum (Fig 6A, B). In support, similar suppression of lysosomal genes by Fxr has been implicated in mouse studies examining Fxr influence in hepatic autophagy. In mice, Fxr trans-represses autophagy-related genes by competing for binding sites with transcriptional activators of these genes, such as CREB (*60, 61*). The binding motif of CREB (“TGACGT”) identified in the mouse study was the second most enriched binding motif near genes repressed by Fxr in zebrafish ileal epithelial cells (Fig S6C) (*61*), suggesting that Fxr may interact with a conserved transcriptional pathway to repress lysosomal functions across these vertebrate lineages. Whereas our data establish roles for zebrafish Fxr on ileocyte and LRE gene expression, we find that Fxr is not required for morphology of the ileal region similar to the observations from *Fxr* knockout mice (*30*). Loss of Fxr function did not overtly affect the relative abundance of ileal epithelial cells (cluster 17) cells, nor the spatial boundaries separating the typical ileocyte region from the adjacent LRE and anterior enterocyte regions (Fig 6C-G). The abundance of LREs also appears unaffected in *fxr* mutants, indicating the observed impacts on LRE gene expression represent altered physiology in those cells (Fig 6A, F). The impacts of *fxr* mutation on ileocyte fate is less clear. Ileocytes are stereotypically defined by their expression of bile salt transport genes, which are markedly reduced in *fxr* mutants as expected (Fig 6A). The differentiation and physiology of the cells that develop in *fxr* mutants within the typical ileocyte region remain unclear, and were not resolved by our scRNA-seq dataset due to the relatively small number of cells located in cluster 17 as well as their substantial heterogeneity (Fig S6). Our results do show that Fxr is involved in tuning distinct transcriptional and physiologic programs of these ileal epithelial cell types while other transcriptional pathways likely determine ileal organization and differentiation. This is consistent with the notion that multiple TFs regulate the same intestinal enterocytes but target distinct cellular processes (*47, 62*).

In contrast to our grasp on Fxr regulation of ileal epithelial cell functions, relatively little is known about the impacts of Fxr on other intestinal cell types. Using scRNA-seq, we show that Fxr exhibits different regulatory effects in anterior enterocytes compared to ileal epithelial cells. Mutation of Fxr led to a significant increase in the abundance of the anterior enterocyte population (Fig 7A, S5A). This is consistent with the observations from intestinal tumorigenesis studies, which show that FXR restricts abnormal stem cell expansion thereby balancing the epithelial proliferative and apoptotic pathways (*30, 63, 64*). It is therefore possible that Fxr similarly affects stem cell dynamics in the zebrafish intestinal epithelium, however such studies await the establishment of markers and tools to study intestinal epithelial stem cells in the zebrafish. Along with the abundance, we also found that the differentiation status of the anterior enterocytes in zebrafish is regulated by Fxr (Fig 7B). Similar roles of Fxr in promoting cell differentiation programs have been reported in other cell types in mammals, including mesenchymal stem cells, adipocytes, and osteoblasts (*65–67*). While the mechanism underlying Fxr’s regulation of cell differentiation remains unclear, we speculate that Fxr may coordinate with Hnf4α to elicit such regulatory effects in zebrafish anterior enterocytes, as Hnf4α binding motif was highly enriched near Fxr-dependent genes (Fig 7C). Indeed, Fxr and Hnf4α can directly interact and cooperatively modulate gene transcription (*68, 69*), and Fxr positively regulates Hnf4α protein levels in mouse liver (*68*). Therefore, it is possible that Fxr increases Hnf4α protein expression or activity to promote enterocyte differentiation in the zebrafish intestine. Nonetheless, our findings reveal novel roles of Fxr in modulating the abundance and differentiation of zebrafish anterior enterocytes.

The molecular and physiologic mechanisms by which Fxr mediates these effects on distinct intestinal epithelial cell types warrant further investigation. Fxr function affects hundreds of genes in the zebrafish (this study) and mouse intestine (*34*), but it remains unclear how many of those are due to primary autonomous roles for Fxr interacting with those gene loci as opposed to secondary systemic effects caused by Fxr mutation. For example, Fxr mutation in the intestine can disrupt endocrine hormone Fgf19 signaling and bile salt homeostasis, therefore producing systemic impacts on energy metabolism, tissue regeneration, and control of inflammation (*70*). Further, Fxr also has critical autonomous roles other organ systems (*71*) which may be impaired in the whole-animal *fxr* mutant zebrafish that we used here. The resulting extra-intestinal and systemic changes may in turn feedback to the intestine to elicit secondary effects on intestinal gene expression and physiology. Tissue-specific and conditional mutant alleles could help distinguish between these different possibilities in the future.

## Materials and Methods

### Zebrafish lines and husbandry

All zebrafish experiments were performed following protocols approved by the Duke University Medical Center Institutional Animal Care and Use Committee (protocol number A115-16-05). Zebrafish stocks were maintained on EK, TL, or a mixed EK/TL background on a 14/10-h light/dark cycle at 28.5 °C in a recirculating system. From 5 dpf to 14 dpf, larval zebrafish were fed Zeigler AP100 larval diet (Pentair, LD50-AQ) twice per day and Skretting Gemma Micro 75 (Bio-Oregon, B5676) powder once per day. From 14 dpf to 28 dpf, larval zebrafish were fed Artemia (Brine Shrimp Direct, BSEACASE) twice per day and Skretting Gemma Micro 75 powder once per day. From 28 dpf to the onset of sexual maturity, Gemma Micro 75 diet was replaced with Gemma Micro 300 (Bio-Oregon, B2809) to feed juvenile zebrafish. After reaching sexual maturity, adult fish were fed Artemia twice per day and a 1:1 mixture of Skretting Gemma Micro 500 and Wean 0.5 (Bio-Oregon, B1473 and B2818) once per day. Male and female adult zebrafish of 3-12 months of age were used for breeding for fish used in this study.

Zebrafish embryos were collected from natural matings and maintained in the corresponding media at a density of <1 larva/mL at 28.5 °C on a 14/10-h light/dark cycle, and were of indeterminate sex. Generation, colonization, maintenance, and sterility test of the germ-free zebrafish larvae were performed as described previously with the exception that an additional 50 μg/mL gentamycin (Sigma, G1264) was supplemented in the antibiotic-containing gnotobiotic zebrafish medium (AB-GZM) (*72*). Conventionally raised zebrafish larvae were maintained in embryo media (0.3% (w/v) crystal sea salt, 0.75 mM CaCO_3_, 0.45 mM NaHCO_3_, methylene blue). The following engineered zebrafish lines were used in this study: *fxr^-10/-10^* (generated in this study), *cyp7a1^-16/-16^* (generated in this study), *Tg(-1.7fabp6:EGFP-pA-cryaa:mCherry)* (generated in this study), *slc10a2^sa2486^* (*25*), *Tg(−4.5fabp2:DsRed)* (*43*), *TgBAC(cldn15la-GFP)* (*31*), *TgBAC(lamp2:RFP)* (*73*), and *Tg(-0.258fabp6-cfos:GFP)* (*17*).

### Genotyping

For genotyping of larval zebrafish, whole larvae were euthanized and placed in PCR tubes containing 50 μL of 50 mM NaOH and denatured at 95 °C for 20 minutes, after which 5 μL of 1M Tris-HCl (pH 8.0) was added. PCR was performed with 2x GoTaq Green master mix using 1 μL template and the corresponding primers (Table S2). For genotyping of adult zebrafish, the tail fin was clipped from anesthetized fish and used to generate the PCR template.

### Construction of mutant zebrafish lines

Mutant zebrafish lines were generated using CRISPR/Cas9 as described previously (*74*). Briefly, the guide RNAs (gRNAs) were designed using the “CRISPRscan” tool (https://www.crisprscan.org/) (*75*) and synthesized using oligo-based *in vitro* transcription method (Table S2). At the one cell stage, wt zebrafish embryos (TL or EK strain) were injected with 1-2 nL of a cocktail consisting of 150 ng/μL of Cas9 mRNA, 120 ng/μL of gRNA, 0.05% phenol red, 120 mM KCl, and 20 mM HEPES (pH 7.0). Injected embryos were screened for mutagenesis with the corresponding primers (Table S2) using Melt Doctor High Resolution Melting Assay (HRMA, ThermoFisher, 4409535) following manufacturer’s specifications. The mutations were further determined through Sanger sequencing of the region encompassing the gRNA targeting sites. The *fxr* mutants were generated through targeted deletion at the exon 4 encoding the DNA binding domain of Fxr. We identified two independent deletion alleles, *fxr^-10/- 10^* and *fxr^-11/-11^* (allele designations *rdu81* and *rdu82* respectively), that each resulted in frameshift mutations and displayed significantly reduced *fxr* mRNA (Fig S1A-B). Likewise, the *cyp7a1* mutants, *cyp7a1^-7-/7^* and *cyp7a1*^-16/-16^ (allele designations *rdu83* and *rdu84* respectively), were generated by targeting the exon 2 encoding the cytochrome P450 domain and were validated via phenotypic assessment and/or qRT-PCR (Fig S1C-F). Only the *fxr^-10/-10^* (*rdu81*) and the *cyp7a1*^- 16/-16^ (*rdu84*) mutants were used in this study.

### Construction of transgenic zebrafish line

The 1.7kb promoter fragment of the *fabp6* gene was PCR amplified from the genomic DNA of wild type Tübingen zebrafish and cloned into p5E-Fse-Asc plasmid (Table S2). The resulting clone (p5E-1.7fabp6), along with the pME-EGFP and p3E-polyA plasmids were further recombined into pDestTol2pACrymCherry through multisite Gateway recombination to generate the pDestTol2-1.7fabp6:EGFPpACrymCherry (*76, 77*). This recombinant plasmid carries two linked fluorescent marker genes, a GFP and a mCherry. The expression of GFP is driven by the 1.7kb *fabp6* promoter fragment and reflects the expression of *fabp6*, whereas the expression of mCherry is driven by the lens marker *cryaa* and serves as a constitutive transgene marker. At the one-cell stage, wt zebrafish embryos (EK strain) were injected with 1-2 nL of a cocktail containing 50 ng/μL pDestTol2-1.7fabp6:EGFPpACrymCherry, 25 ng/μL transposase mRNA, 0.3% phenol red and 1x Tango buffer (ThermoFisher, BY5). Two mosaic germline founders were identified, raised to adulthood, and screened to isolate lines with the transgene inserted at a single locus. Stable *Tg(-1.7fabp6:EGFP-pA- cryaa:mCherry)* (allele designations *rdu80*) lines were generated by outcross the founder to wt EK for at least three generations (abbreviated as *Tg(-1.7fabp6:GFP)* in the rest of the article). This *Tg(-1.7fabp6:GFP)* reporter line displayed a pattern of GFP expression in the ileocyte region and the LRE region similar to our previous transgenic line *Tg(-0.258fabp6 -cfos:GFP)* (*17*) which expresses GFP under control of a smaller 258bp *fabp6* promoter region and a mouse *Cfos* minimal promoter. Compared to that line, the new *Tg(-1.7fabp6:GFP)* line using the larger 1.7kb *fabp6* promoter region expresses GFP more distally into the LRE region and also in rare cells within the anterior regions of the intestine.

### Quantitative RT-PCR analysis

RNA was isolated from samples using TRIzol (ThermoFisher, 15596026), DNase-treated using TURBO™ DNase (ThermoFisher, AM2238), and reverse transcribed using iScript cDNA synthesis kit (Bio-Rad, 1708891) following manufacturer’s specifications. Quantitative PCR was performed with gene-specific primers (Table S2) and SYBR Green PCR Master Mix with ROX (PerfeCta, Quanta Bio) on an Applied Biosystems StepOnePlus™ Real-Time PCR System. Data were analyzed with the ΔΔCt method. For whole larvae samples, 6-7 dpf larvae were collected for RNA isolation (15-30 larvae/replicate; 4-8 replicates/condition). For larval digestive tissue samples, 6-7 dpf larval zebrafish digestive tracts were dissected under a stereomicroscope and pooled for RNA isolation (25-35 guts/replicate; 4-6 replicates/condition). For adult digestive tissue samples, 3-month-old gender and size matched adult zebrafish livers or guts were used (1 gut or liver/replicate; 5-8 replicates/condition).

### *In vivo* imaging and densitometry

To quantify the GFP fluorescence of the *Tg(-1.7fabp6:GFP)* lines, live zebrafish larvae were anesthetized, embedded in 3% methylcellulose (w/v in GZM), and imaged using a Leica M205 FA stereomicroscope with identical exposure time and magnification in the same experiment. GFP densitometry analysis was performed using Fiji software (*78*). For each experiment, the areas of interest were selected using the shape tools, recorded using the ROI manager, and applied to all images. The background was calculated as the average fluorescence from 3-5 non-transgenic siblings of the transgenic zebrafish lines from the same experiment and was subtracted from all images using the threshold tools. The mean fluorescence intensity values of each image were determined and plotted using GraphPad Prism software.

### Bile salts collection in zebrafish

Twenty wild-type adult zebrafish of 6-9-month-old from 4 different stocks were starved for 48 h to eliminate the potential contribution of exogenous bile salt from the zebrafish diet on zebrafish *de novo* bile salts. The gallbladders were dissected using autoclaved forceps and immediately placed in 1 mL of pre-chilled isopropanol. The suspension was vortexed and centrifuged (13,000 rpm for 10 min), and the supernatant was evaporated under a stream of nitrogen at room temperature. The residues were resuspended in 1 mL of 100% methanol. For thin layer chromatography (TLC), 20 µ L of the methanol extract was spotted on to a silica TLC plate. The butanol: acetic acid: water (85:10:15, BAW) mobile phase system was used to separate the bile components. Additionally, a diluted (1:100) sample was subjected for LC/MS analysis. For bile salt analysis of zebrafish intestinal contents, 10 pooled wt adult zebrafish intestines were subjected to the same procedures as gallbladders above and diluted (1:100) for LC/MS.

### Flash column chromatography of crude carp bile

Asian grass carp gallbladders (n= 3) were collected from a local supermarket in Champaign, IL. Bile was collected from each gallbladder and pooled for extraction (45 mL). Crude bile was extracted using 9x isopropanol, and the isopropanol-soluble portion was collected for analysis. The isopropanol layer was concentrated to approximately 20 mL under nitrogen. Diluted (1:100) crude bile samples were used for TLC analysis with the BAW mobile phase. Purification of carp bile acids and alcohols was performed using flash column chromatography as described previously (*23*). The flash column (80 cm x 2 cm; 100 mL) was packed 2/3 full with 40 µ M silica gel. It was assembled using chloroform: methanol (80:20; v/v) mobile phase. The concentrated isopropanol-bile mixture was placed on top of the packed silica for purification. The eluates of crude bile were collected in 50 mL fractions using a gradient of chloroform:methanol (80:20; 500 mL, 75:25; 500 mL, 70:30; 1000 mL, 65:35; 500 mL). The fractions were evaporated under nitrogen and resuspended in 100% methanol. A dilute sample of each fraction was spotted (30 µ L) and examined on a TLC plate using BAW mobile phase.

### Select fractions were chosen for LC/MS analysis

TLC visualization and extraction of bile compounds from TLC Zebrafish and carp bile in methanol were examined using silica gel TLC plate (JT Baker, JT4449-4). Two mobile phases were used to separate bile alcohols and bile acids. BAW mobile phase consisted of butanol: acetic acid: water (85:10:15) mobile phase. Solvent 25 mobile phase used was n-propanol: isoamyl acetate: acetic acid: water (4:3:2:1). Plates were sprayed with 10% phosphomolybdic acid (w/v) in ethanol and plates were baked at 100 °C for 10 min. To extract bile compounds from the TLC plate, silica from replicate plates was extracted twice with 3 mL butanol and 3 mL water. The butanol layer was removed after each extraction, combined, and evaporated under nitrogen gas.

### Extraction of carp intestinal contents

Whole intestines were removed from Asian carp and collected in 50 mL conical tubes. The contents were placed in a -80 °C freezer overnight and lyophilized to remove all liquid. For LC/MS analysis, dry intestinal contents (0.14 g) were resuspended in 1 mL of 90% ethanol and sonicated for 30 min to completely dissolve soluble compounds. Furthermore, the intestinal content was centrifuged (10,000 rpm for 15 min) and the supernatant was filtered (0.45 µ m) to remove additional precipitates. Diluted samples (1:100) of the filtered supernatant were spotted (30 µ L) on to a TLC using BAW mobile phase and also injected on to LC/MS in untargeted full scan mode to analyze metabolites.

### NMR analysis of purified zebrafish bile alcohol

Pure flash column chromatography fractions and TLC spots matching the Rf value for 5αCS were validated using mass spectrometry in negative ion mode. 1 mg of pure bile alcohol in methanol was used on a Waters SynaptG2-Si ESI MS. The MS data was analyzed using Waters MassLynx 4.1 software. Additionally, a 4 mg sample of the evaporated bile alcohol was resuspended in 750 µ L of deuterated methanol and analyzed by nuclear magnetic resonance spectroscopy using an Agilent 600 MHz with a 14.1 Tesla 54 mm bore Agilent Premium Compact Shield Superconducting Magnet. Data was visualized at the University of Illinois using MNova.

### Liquid chromatography/mass spectrometry (LC/MS)

LC/MS for all samples was performed using a Waters Aquity UPLC coupled with a Waters Synapt G2-Si ESI MS. Chromatography was performed using a Waters Cortecs UPLC C18 column (1.6 µ m particle size) (2.5 mm x 50 mm) with a column temperature of 40°C. Samples were injected at 1 µ L. Solvent A consisted of 95% water, 5% acetonitrile, and 0.1% formic acid. Solvent B consisted of 95% acetonitrile, 5% water, and 0.1% formic acid. The initial mobile phase was 90% Solvent A, 10% Solvent B and increased linearly until the gradient reached 50% Solvent A and 50% Solvent B at 7.5 min. Solvent B was increased linearly again until it was briefly 100% at 8.0 min until returning to the initial mobile phase (90% Solvent A, 10% Solvent B) over the next 2 min. The total run was 10 min with a flow rate of 10 µ L/min. MS was performed in negative ion mode. Nebulizer gas pressure was maintained at 400 °C and gas flow was 800 L/hour. The capillary voltage was set at 2,000 V in negative mode. MassLynx was used to analyze chromatographs and mass spectrometry data. The limit of detection (LOD) was defined as a 3:1 signal to noise ratio using the LC peak data. The limit of quantification was defined as the 10:1 signal to noise ratio using the LC peak data. A mixture containing 10 µ M of the following bile standards were injected onto LC/MS for analysis: D4-Glycocholic acid (Internal Standard), TCA, 5αCS, and allocholic acid (ACA). The LC/MS method was validated once a single peak for each compound was identified with the respective m/z value in negative mode.

To test the bile salt metabolism activity of complex zebrafish microbiota, the contents from the dissected intestines of 6 wt adult zebrafish from 4 different stocks were pooled into 4 samples as representatives of distinct zebrafish microbial communities. Each sample was homogenized in 500 µ L PBS with 1 mM DTT. The resulting intestinal homogenate was split into both aerobic and anaerobic vials containing modified TSB (1:10 dilution). Aerobic cultures were incubated with 200 rpm shaking while anaerobic cultures were incubated statically. Both aerobic and anaerobic cultures were incubated at 30 °C for 24 h, after which they were subcultured (1:10 dilution) into different substrate testing media and allowed to grow at 30 °C for an additional 48 h before being subjected to solid phase extraction.

To test the bile salt metabolism activity of individual microbial strains, *Pseudomonas* sp. ZWU0006, *Acinetobacter* sp. ZOR0008, *Shewanella* sp. ZOR0012, *Exiguobacterium acetylicum* ZWU0009, and *Chryseobacterium* sp. ZOR0023 were grown at 30 °C for 24 h and were subcultured (1:10 dilution) into different substrate testing media, respectively. The subcultures were grown under the same condition for an additional 48 h and were subjected to solid phase extraction.

### Solid phase extraction of bacterial culture

Culture medium (1 mL) containing 50 µ M bile salt substrate was used for further SPE. Once grown, the culture was centrifuged (10,000 rpm for 5mins) to remove bacterial cells and conditioned medium was removed. A 10 µ M spike of D4-GCA internal standard was added to each sample before SPE. Waters tC18 vacuum cartridges (3 mL reservoir, 500 mg sorbent) were used for SPE. The method was adapted from Abdel-Khalik, et al as follows (*79*). Cartridges were preconditioned with 100% hexanes (6 mL), 100% acetone (3 mL), 100% methanol (6 mL), and water adjusted to pH 3.0 (6 mL). Conditioned medium was adjusted to pH 3.0, applied to the cartridge, and pulled through dropwise using a vacuum chamber. The cartridge was washed with water adjusted to pH 3.0 (6 mL) and allowed to air dry for 30 min before being washed with 3 mL of 40% methanol. The 40% methanol fraction was tested on TLC to ensure no substrates were being washed off of the column. Products were eluted using 3 mL of 100% methanol. Final eluates were evaporated under a stream of nitrogen and resuspended in 200 µ L of 100% methanol for analysis on TLC (using solvent 25) or LC/MS.

### Serial bile salt exposure

For serial bile salt exposure in conventionally raised larvae, embryos were collected from natural matings between *cyp7a1*^+/-16^ and *cyp7a1*^+/-16^; *Tg(-1.7fabp6:GFP)* and incubated in GZM at 28.5 °C. At 3 dpf, larvae were randomly assigned into untreated group or groups treated with either 1 mM TCA or CA in GZM in 6-well plates. The density of larvae in each well is maintained as 10 larvae in 10 mL media and the media in each well were changed daily (80% v/v) with fresh GZM or GZM supplemented with either 1 mM TCA or CA. At 7 dpf, larvae were sorted for mCherry under a fluorescence microscope, after which positive larvae were subjected to *in vivo* imaging and genotyping. Serial bile salt exposure in GF larvae was performed similarly except that GF larvae were maintained in T25 tissue flasks and that sterile TCA or CA was used for treatment.

### Fluorescence-activated cell sorting (FACS)

Approximately 600-700 6 dpf *TgBAC(cldn15la-GFP)* zebrafish larvae of the *fxr^+/+^* and the *fxr^-/-^* genotypes were collected for the FACS experiment, respectively. The parental zebrafish used to generate the wt or the mutant embryos were stock-matched siblings from heterozygous incrosses. Dissociation of the larvae was performed as previously described (*80*), after which the *fxr*^+/+^; *TgBAC(cldn15la-GFP)* and *fxr*^-/-^; *TgBAC(cldn15la-GFP)* cells were immediately subjected to FACS at the Duke Cancer Institute Flow Cytometry Shared Resource and were sorted side by side with two identical Beckman Coulter Astrios instruments. Non-transgenic and single transgenic controls (pools of 50 fish/genotype) were prepared as above and used for gating and compensation. Approximately 120 k GFP positive 7-AAD negative cells per genotype were collected in 1.5 mL of DMEM/F12 supplemented with 10% heat-inactivated FBS and 10 μM Y- 27632 ROCK1 inhibitor and were immediately subjected to the downstream experiments.

### Single-cell RNA sequencing

Each single-cell RNA sequencing library was generated from 10,000 FACS sorted *TgBAC(cldn15la-GFP)* IECs of the indicated genotype following the 10x Genomics Single-cell 3’ protocol by the Duke Molecular Genomics Core. The sequencing ready libraries were cleaned with both Silane Dynabeads and SPRI beads, and quality controlled for size distribution and yield with the Agilent D5000 screenTape assays using the Agilent 4200 TapeStation system. Illumina P5 and P7 sequences, a sample index, and TruSeq read 2 primer sequence were ligated for Illumina bridge amplification. Sequence was generated using paired-end sequencing on the Novaseq SP flow cell sequencing platform at a minimum of 40 k reads/cell.

Cell barcodes and unique molecular identifier (UMI) barcodes were demultiplexed and reads were aligned to the reference genome, *danRer11*, following the CellRanger pipeline recommended by 10X Genomics. For quality control, we first performed UMI filtering by only including UMIs with <3000 detected genes. Next, we removed low quality cells which we define as cells that contain >25% transcript counts derived from mitochondrial genes. Further, we removed the putative doublets by excluding cells that contain more than 30,000 UMIs. Through these steps, a total of 2,625 low-quality or potential doublet cells were removed, after which 9,918 cells passed the requirement, including 4,710 cells from *fxr* wt and 5,208 cells from *fxr* mutant samples. The genotype of *fxr* wt and mutant samples was confirmed by visualization of reads spanning the -10/-10 lesion (Fig. S4A).

Clustering and statistical analysis of the single-cell RNA-sequencing data was performed using the R package Seurat (version 3.1). Count matrices from both the *fxr* wt and mutant libraries were log-normalized and highly variable genes were found in each library using the FindVariableFeatures() function. Afterwards, these data were integrated together using the wt library as the reference dataset through the FindIntegrationAnchors (dims = 1:35) and IntegrateData (dims = 1:35) functions. The integrated expression matrix was then re-normalized using the NormalizeData() function for visualization purposes. To mitigate the effects of unwanted sources of cell-to-cell variation in the integrated dataset, we used the ScaleData() function prior to running a principal component analysis. Jackstraw analysis revealed that the first 54 principal components significantly accounted for the variation in our data, and were thus used as input to the FindClusters() function with the resolution parameter set to 0.82. Using the shared nearest neighbor algorithm (SNN) within the FindNeighbors() function, cells were grouped into 27 distinct clusters and were visualized by uniform manifold approximation and projection (UMAP), which reduces the information captured in the selected significant principal components to two dimensions. The UMAP visualization was generated using the RunUMAP() function with the “n_neighbors” parameter set to 30.

To resolve putative distinct functional cell types in cluster 17 cells in *fxr* wt zebrafish, we performed sub-clustering of the cluster 17 using a similar strategy as described above with the exception that we used 8 principal components following JackStraw analysis and a resolution of 0.5 in the FindClusters() function. This resulted in two sub-clusters: 17_0 and 17_1 (Fig S6D).

To identify marker genes of *fxr* wt cells in each cluster, we used two methods with different stringency standards. First, we employed the FindAllMarkers() function using a Wilcox Rank Sum Test to determine genes that are significantly upregulated in each cluster compared to all other clusters combined as one group. These genes were further filtered based on an adjusted p-value below 0.05 and an absolute log_10_ fold-change value over 0.25, resulting in a set of marker genes that we designated as “cluster markers” (Dataset 2). Second, we performed pairwise comparisons between the cluster of interest and each and every other clusters using FindMarkers() function and only selected genes that showed higher expression, defined as an absolute log_10_ fold-change value over 0.25, in the cluster of interest in all comparisons. This pairwise comparison-based filtering step resulted in a set of more stringent marker genes, designated as “cluster-enriched markers”, that represented the most highly expressed genes in the cluster of interest (Dataset 3). The expression and the distribution of relevant cluster markers or any gene of interest were visualized using FeaturePlot(), DotPlot(), and VlnPlot() functions.

To identify genes that were differentially expressed between the *fxr* wt and mutant cells in each cluster, we used the FindMarkers() function using a Wilcox Rank Sum Test. Differentially expressed genes were arbitrarily defined as those that showed an absolute log_10_ fold-change value over 0.25.

### Transcription factor binding motif enrichment analysis

We used FAIRE-Seq data from adult zebrafish intestinal epithelium (*17*) to identify accessible chromatin regions at genes that are differentially regulated in either cluster 17 or 4. Using GALAXY, each FAIRE-Seq peak was associated with the nearest gene, including its surrounding regulatory regions (including 10kb from the gene transcription start site, the gene body, and 10kb from transcription termination sequence). We generated a BED file containing this information for every gene, that could be filtered based on gene symbol identifier later based on whether or not a particular gene was differentially expressed in clusters 17 or 4. To identify enriched transcription factor binding sites, we used “findMotifsGenome.pl” function of the HOMER software (http://homer.ucsd.edu/homer/) with foreground and background set of genomic coordinates. Specifically, genes that were differentially expressed between *fxr* wt and mutant cells in the cluster of interest were used as the foreground, and the ones that were not differentially expressed in the cluster of interest but exhibited expression in at least one of the IEC clusters were used as background.

### Statistical Analysis

For the scRNA-seq experiment, statistical analyses for determination of the cluster markers, cluster-enriched markers, and differentially expressed genes of each clusters were calculated using the FindMarkers() function of the Seurat package in R with a Wilcox Rank Sum Test. For all other experiments, statistical analysis was performed using unpaired t-test, or one-way or two-way ANOVA with Turkey’s multiple comparisons test with GraphPad Prism. A P<0.05 was defined as statistically significant.

## Acknowledgments

**General:** We thank Dr. Jieun Esther Park, Dr. Daniel Levic, and Laura Childers for their technical assistance. We extend our gratitude to Dr. Furong Sun, director of the Illinois Mass Spectrometry Core for providing assistance with LC/MS analysis. We would like to acknowledge the assistance of the Duke Molecular Physiology Institute Molecular Genomics Core for the generation of scRNA-seq data.

## Author contributions

J.W., J.M.R., and J.F.R. designed research; J.W. and A.V. performed research; G.K., C. K., and J.L.C contributed new reagents/analytic tools; J.W., G.P.M., A.V., H.L.D., C.R.L., T.C., G.K., J.M.R, and J.F.R. analyzed data; and J.W., G.P.M., A.V., H.L.D., C.R.L., J.M.R., and J.F.R. drafted and revised paper.

## Data and materials availability

All data needed to evaluate the conclusions in the paper are present in the paper and/or the Supplementary Materials.

## SUPPLEMENTARY MATERIALS

### Supplementary Results

#### Annotation of single-cell RNA-seq data identifies cell types within the larval zebrafish intestinal epithelium

We generated scRNA-seq data on GFP+ cells sorted from wild-type (wt) and *fxr^-^*^/-^ *TgBAC(cldn15la-GFP)* zebrafish larvae at 6 dpf. The *TgBAC(cldn15la-GFP)* transgenic line was chosen for this study because it is reported to express GFP throughout the intestinal epithelium (*31*). After quality control filtering, a total of 9,918 cells from both genotypes combined were subjected to unsupervised clustering using the Seurat R package yielding a total of 27 clusters (Fig 5A). As expected, the vast majority of cells and clusters expressed *cldn15la* and displayed gene expression patterns consistent with known gut epithelial cell types. Below we provide a working annotation of each cluster as a resource for the field. For this annotation, we focus on data derived from wild-type zebrafish intestinal epithelial cells. Our general approach was to study the genes we found to be enriched in each cluster or set of related clusters, along with UMAP visualization of the gene expression patterns in the current scRNA-seq dataset, the primary literature, and gene expression patterns reported on ZFIN. Due to limited previous studies of the zebrafish digestive tract, we acknowledge that some of these annotations may be found to be inaccurate in future studies. In addition to the annotation below, we provide average expression of genes in individual cell clusters (Data set 1), cluster markers of individual cell clusters identified (Data set 2), and cluster-enriched markers of individual cell clusters identified (Data set 3).

*Enterocytes:* The principal function of the intestinal epithelium is absorption of dietary nutrients, provided primarily by enterocytes. As described in further detail in the Results section, we identified clusters 4, 9, 16, and 17 as enterocytes based on their expression of known enterocyte markers. We tentatively assigned these clusters as enterocytes in the anterior intestine (also called the intestinal bulb or segment 1; clusters 4 and 16), ileal/mid-intestine (also called segment 2; cluster 17), and distal intestine/cloaca/pronephric duct (also called segment 3; cluster 9). Clusters 4 and 16 were both enriched for known anterior enterocyte markers involved in metabolism and transport of lipids (e.g., *rbp2a*, *fabp1b.1*, *apoa1a*, *apoa4b.2.1*, *apobb.1*, *scarb1*), chitin (e.g., *chia.1*, *chia.2*), and other genes not previously implicated in anterior enterocyte biology (e.g., *mbl2*, *rida*, *gcshb*). Despite these commonalities, these two anterior enterocyte clusters also displayed interesting differences. Metascape analysis of enriched markers in cluster 4 revealed overrepresentation in biological processes such as oxidation-reduction (e.g., *fads2*, *agmo*, *aoc1*), gluconeogenesis (*fbp1b*, *pck1*), and metabolism of lipids and other nutrients (e.g., *apoa1b*, *apoea*, *pla2g12b*, *elovl2*, *slc26a3*.2, *slc34a2a*, *slc37a4a*, *slc6a19b*) (Fig S8). In contrast, markers enriched in cluster 16 include those involved in chromatin organization (e.g., *hist2h3c*, *si:ch211-113a14.18*, *zgc:110425*, *zgc:173585*), and cell cycle and chromosomal segregation (e.g., *mki67*, *cks1b*, *cks2*, *nuf2*, *aurkb*, *cdca8, birc5a*, *kif20bb*, *plk1*, *spc25*, *top2a*, *ube2c*) (Fig S8). Based on these observations, we operationally define cluster 4 as differentiated anterior intestinal enterocytes and cluster 16 as proliferative anterior intestinal enterocytes.

Whereas clusters 4 and 16 appear to represent anterior enterocytes, clusters 17 and 9 appear to represent enterocytes from the mid and distal regions of the intestine. As described in detail in the Results, cluster 17 was identified as mid-intestinal enterocytes which include ileocytes and lysosome-rich enterocytes (LREs) (see Figs 6 and S6, Table S1, Dataset 1-2). Metascape analysis of cluster 17 revealed enrichment for bile acid and bile salt transport among other functions (Fig S8), Cluster 9 expresses known distal intestinal epithelial markers like *saa* (*43*), *irg1l* (*81*), *hoxa13a* and *evx1.1* (*82*). This cluster also expresses *ponzr1* and *slc9a3.2* which are expressed not only in the intestine but also the pronephric duct, tubules, and nephrons (*83, 84*). This raises the possibility that this cluster may include cells from the pronephric duct or tubule. However, we think this is unlikely since cluster 9 does not express several known markers of the pronephric duct and tubule epithelium like *atp1a1a.5* (*84*) and *cdh17*, *pax2a*, *slc4a2a* (*85*). Metascape analysis for cluster 9 was not particularly informative (Fig S8), but based on the above information we predict that cluster 9 represents enterocytes from the distal intestine and cloaca.

Absorptive enterocytes are typically the most common epithelial cell type in the intestine, so we were surprised that the relative abundance of cells in clusters 4, 16, 17, and 9 was comparable to other cell types discussed below. We expect that these relative cell abundances are inaccurate and infer that our chosen methods for cell dissociation, sorting, processing, and/or data quality may have preferentially led to a reduced representation of enterocytes in this dataset. However, we think these four enterocyte clusters do represent the major enterocyte populations in the larval zebrafish intestine. Although cluster 16 appears to be a proliferative enterocyte population, additional studies are needed to identify absorptive lineage precursors that give rise to these different differentiated enterocyte types.

*Goblet cells:* Mucus provides an important physical and chemical barrier along the gut, and is primarily produced by secretory goblet cells. Anterior gradient 2 (*agr2*) is expressed in goblet cells within the zebrafish larval gut, as well as the pharynx and esophagus (86, 87), and Agr2 is required for goblet cell development and mucin production in zebrafish and mice (87–89). SAM Pointed Domain Containing ETS Transcription Factor (*Spdef*) is expressed by and promotes differentiation of goblet cells and Paneth cells in the mouse intestine (90, 91), however fishes are not thought to develop Paneth cells. We found that clusters 3, 13, and 14 all express high levels of agr2, two of which (clusters 3 and 14) also displayed elevated expression of spdef. An additional shared marker for these three clusters was *galnt7* which is involved in mucin-type O-linked protein glycosylation. Goblet cells do produce mucins at this stage (41), but the mucin genes detected in our scRNA-seq dataset did not show enrichment in these three clusters. Cluster 13 displayed elevated expression of POU Class 2 Homeobox 3 (*pou2f3*), which is a lineage-specifying transcription factor for tuft cells in mice (92), as well as Sprouty 2 (*spry2*) which is regulates differentiation of goblet and tuft cells in mice (93). Interestingly, cells in cluster 13 also displayed elevated expression of genes known to be involved in the development of antigen-sampling microfold cells (M-cells) in the mammalian intestine including SRY-Box Transcription Factor 8 (*sox8b*) (94), TNF Receptor Superfamily Member 11a (*tnfrsf11a*; also known as RANK) (95), and its conserved paralog *tnfrsf11b* (also known as Osteoprotegerin) which acts as a decoy receptor for RANK ligand (96). Tuft cells and M-cells are considered to be distinct differentiated cell types in the mammalian gut, but these zebrafish cells in cluster 13 appear to display conserved markers of both cell types. Based on these criteria, we annotated these three clusters as goblet cells, with the possibility that cluster 13 represents zebrafish tuft cells or M-cells.

*Enteroendocrine cells:* The other major secretory cell lineage in the intestine is enteroendocrine cells (EECs) which serve as specialized sensory cells that respond to luminal and basolateral stimuli to release specific hormones and neurotransmitters. We identified five clusters of cells (clusters 8, 11, 12, 21, and 22) as putative EEC subtypes based on their shared expression of known EEC markers *neurod1* (97), *pax6b* (98), and *nkx2.2a* (99). Those same five clusters expressed other known markers of EECs such as *pcsk1*, *gck*, and the Secretogranins *scg2b*, *scg3*, and *scg5*. GO Term Enrichment analysis of each of these clusters revealed pathways associated with protein processing and transport, with some specific pathways linked to neurotransmitters and neuropeptides. As expected, each of these EEC subtypes was characterized by distinct gene expression signatures for transcription factors, hormones, and neurotransmitters. For example, EEC subtypes displayed distinct expression patterns for known EEC transcription factor homologs neurog3 and pax4 (cluster 11), isl1 (clusters 8, 21, and 22), pdx1 (cluster 22), and lmx1ba (cluster 12) (100, 101). Genes involved in production of hormones, neurotransmitters, and other signaling molecules also showed elevated expression in clusters 8 (*ghrl, pyyb, sst2, mlnl*), 11 (*trhra*), 12 (*adcyap1a, bdnf, hbegfb, il22, nmu, penka, tph1b, trpa1b*), 21 (*calca*), and 22 (*gcga, insl5b, tac3a, galn, btc, vipb*). Other hormones and neurotransmitters were expressed in multiple EEC clusters, such as *ccka* and *nmbb* (clusters 11 and 12) and *insl5a* (clusters 21 and 22). Based on these gene expression patterns, potential homologies to known mammalian EEC subtypes were suggested. For example, cluster 11 may represent early differentiating EECs due to their expression of transcription factors involved in early EEC differentiation *neurog3* and *pax4*, and their relative paucity of hormone or neurotransmitter expression. Like cluster 8 EECs, motilin-producing EECs (M-cells) in the mammalian gut also express Ghrl, Sst, and Mln. Like cluster 22, L-cells in the mammalian gut also express the proglucagon gene Gcg (the precursor to GLP-1 and GLP-2) as well as Insl5 (102, 103). Finally, zebrafish EEC cluster 12 and mammalian enterochromaffin cells (ECs) express respective homologs of Tph1 and Trpa1, as well as other shared markers Rab3c and Ddc (44, 104). We anticipate that some of these individual EEC clusters may represent multiple EEC cell types which could be further resolved by future studies. Together, these data reveal interesting functional diversity among EEC subtypes in larval zebrafish, including potential subtype conservation that has been conserved since the last common ancestor with mammals.

*Putative secretory precursors:* Studies in the mammalian intestine identify Math1/Atoh1, Dll1, and Gfi1 as markers of secretory lineage precursors (105). In our dataset, zebrafish *math1/atoh1* and *dll1* homologs were not detected, whereas *gfi1b* was expressed in a subset of cells in cluster 13 (goblet/tuft cells) and *gfi1aa* was expressed in a small fraction of cells across multiple secretory and absorptive cell clusters. Despite this paucity of clear conserved markers, two clusters presented as potential precursors of secretory cell lineages based on their location in the UMAP plot and their gene expression patterns. Clusters 5 and 6 both expressed agr2 though at lower levels than the goblet cell clusters. Cluster 5 appeared to be either EEC precursors or perhaps another differentiated EEC subtype based on its strong expression of pyyb (similar to cluster 8 EECs), and expression of *pax4*, *neurod1*, *scg2b*, *scg3*, *scg5*, *insl5a*, and *mlnl* in a subset of cells. Located adjacent to cluster 5 and the goblet cell clusters in the UMAP plot, cluster 6 was enriched for transcripts involved in the H/ACA small nucleolar ribonucleoprotein (H/ACA snoRNP) complex including nhp2, nop10, dkc1, and gar1 (106). Additional cluster 6 markers include other ribonucleoproteins *nop58, snu13b, hnrnpabb, and npm1a*, RNA processing and splicing factors like *ssb, trmt61a, ddx39ab, ddx24, and c1qbp*, and other markers like the nucleolar G-protein *gnl3* and the dihydrolipoamide dehydrogenase *dld*. Although the precise identity of cluster 6 remains unclear, their location on the UMAP plot combined with their enrichment for these markers for RNA processing and ribosomal biogenesis suggests they are likely secretory precursors at a stage of differentiation marked by elevated transcription and translation. Since the lineage relationships between intestinal epithelial cells and markers for intestinal epithelial stem cells remain unknown in the zebrafish, it is also possible that cells within cluster 6 represent common precursors of both secretory and absorptive lineages.

*Ionocytes:* Electrolyte secretion and ion balance are important aspects of epithelial biology in the intestine and other organs (107). Epithelial cells specialized to perform these functions, typically called ionocytes (sometimes called chloride cells), can be found in diverse organ systems including the human airway, frog epidermis, and fish epidermis and gills (108–112). The chloride and bicarbonate ion channel Cftr is known to be a marker of ionocytes in the mouse and human airway as well as fish. In the small intestine of human and rat, Cftr is expressed at high levels in a distinct subset of cells (113, 114) though their function remains unclear. We found that a distinct group of three clusters 0, 2, and 15 were enriched for cftr, as well as other markers conserved with mammalian ionocytes like *tmem51a* and *stap2b* (108, 109). These three clusters also express the carbonic anhydrases *ca2* and *ca4b*, which may function to produce bicarbonate for subsequent Cftr-mediated transport across the plasma membrane. A recent study identified a new population of epithelial cells in the human colon predicted to be involved in electrolyte transportation marked by the proton-conducting ion channel Otopetrin 2 (OTOP2), the calcium-sensitive chloride channel Bestrophin 4 (BEST4), and the paracrine hormone uroguanlylin (33). Strikingly, we found that clusters 0, 2, and 15 express otop2, best4, and the predicted receptors for uroguanylin (*gucy2c*) and atrial and brain natriuretic peptides (*npr1a*). These data strongly suggest that these cells are specialized for regulated secretion of bicarbonate and other ions to control mucus secretion (115) and perhaps other processes. We therefore conclude that these clusters together represent intestinal ionocytes, with similarities to mammalian BEST4/OTOP2 cells. Interestingly, these three ionocyte clusters were also enriched for notch2 and the Notch-responsive gene her6/hes1. In the mammalian intestine, *Her6/Hes1* and Notch2 are expressed in the crypt in stem cells and absorptive progenitors (116, 117). In accord, we previously identified a cis-regulatory element at zebrafish her6/hes1 that drove expression in IECs near the base of intestinal folds that also activated a Notch reporter (17), in accord with previous studies of zebrafish Hes1 localization (118). These findings suggest a previously unappreciated relationship between Notch signaling and intestinal ionocytes in zebrafish. The significance of other genes enriched in all three ionocyte clusters (e.g., *cfd, osr2, syt7b, fgfr4, abcc12, prdx1, hmox1a, syt7b, si:dkey-190j3.2, tmtops2b*) or individual clusters (e.g., *hsp70.1, hsp70.2, hsp70.3, hsp70l, and hsp90aa1.2* for cluster 15; *osr2, gsto1, scpp8, and tpmt.1* for cluster 2; *epb41a and MPRIP* for cluster 0) remains unclear.

*Foregut:* The transcription factor Sox2 in birds and mammals is expressed in foregut regions including pharynx, esophagus, and stomach (119). This expression domain is conserved in the stomachless zebrafish, with *sox2* expressed in pharyngeal and esophageal epithelium (120). This foregut marker was only expressed in the distinct group of clusters 1, 7 and 10. Annotation of these individual clusters was more difficult than others due to a relatively low number of genes enriched specifically in those individual clusters. However, there were several genes enriched in all three clusters that support that these clusters represent epithelial cells from foregut regions such as pharynx and esophagus. For example, all three clusters expressed known markers of the pharyngeal epithelium such as *ca15* (121), *aqp3a* (122), *ptgs2b*, and *ptgs2a* (123). These three clusters were enriched for other known pharyngeal markers that were also expressed at lower levels in other non-pharyngeal clusters such as *wu:fb18f06* (124) and *sult6b1* (125). Notably, these three clusters and the anterior enterocyte clusters 4 and 16 shared expression of *fabp2*, which is typically considered to be a marker of anterior intestinal enterocytes (43). However, the absence of other known anterior intestinal enterocyte markers (e.g., *rbp2a, fabp1b.1, apoa1a;* see above) in clusters 1, 7, and 10 lead us to conclude those three clusters do not represent anterior intestinal enterocytes. These three clusters were also enriched for other genes without available WISH data including *si:ch73-288o11.5, sytl4, si:dkey-74k8.3, icn, ktn1, ARF5* (1 of many), *cdx1a, vill, vtcn1, and gstp2*, which may represent new markers for these cell types. We observed that several of the genes enriched in clusters 1 and 10 were also enriched in cluster 9 which we annotate as enterocytes from the distal intestine and cloaca. These include prdm1a which is a known marker of the cloaca (126–128), the pharyngeal marker bcam (129), and other genes that lack WISH data (*CABZ01020840.1, BX908782.2, efna1b, lect2l, si:cabz01007794.1, zgc:113314*). It is notable also that agr2, which is expressed most highly in goblet cell clusters 3, 13, and 14, is also expressed at lower levels in foregut clusters 1, 7, and 10 as well as distal enterocyte cluster 9. As cells lining the entrance and exit of the intestinal tract, it is tempting to speculate these genes may represent unknown shared physiologic functions in those cells. We were unable to functionally distinguish these three individual clusters due to paucity of specifically enriched genes. Cluster 10 specifically expressed several interesting genes including the pharyngeal markers *capn2a* (130) and *muc5.3* (131), mucin synthesis enzyme *gcnt3*, a cysteine-rich natrin-1-like venom protein (CABZ01068499.1), O-glycan processing enzyme *si:dkey-202e17.1* as well as *si:dkey-248g15.3* and *arrdc2*. Genes enriched in cluster 10 but also expressed appreciably by other clusters includes the pharyngeal marker rhpn2 (125), as well as *aplp2, c7b, and tnfrsf9a*. In contrast, clusters 1 and 7 revealed very few genes that were specific to these clusters with the nearest examples being *neu3.3 and si:ch211-284d12.3* in cluster 7, and *CU467905.1 and stoml3b* in cluster 1. Based on these observations, we annotate clusters 1, 7, and 10 as foregut epithelial cells. Since publicly available WISH data provide more information about pharyngeal gene expression compared to esophagus, we are unable to confidently annotate these three clusters to specific regions of the foregut at this time. Cluster 10 seems likely to be mucus producing cells but the specific identities of clusters 1 and 7 remain unresolved. We note that known markers of the oral epithelium such as *evplb* (124), *brpf1* (132), and *barx2* (133) were low or absent in our dataset. Therefore, while we annotate clusters 1, 7, and 10 as foregut epithelial cells, we do not exclude that the oral cavity or other foregut regions include cell types not captured in our dataset here.

*Epidermis:* Analysis of genes enriched in clusters 23 and 26 suggest they are integumentary cells. Cluster 23 expresses *tp63*, a known marker of the basal layer of the epidermis (134–136), as well as other genes with known expression in the epidermis such as *ecrg4b* (137), *krt5*, *mmp9* (138), *zgc:101810* (122), *col4a5, col4a6* (139), and *col17a1a* (140). Another gene specifically enriched in cluster 23 specific for cluster 23, cldni, is reported to be expressed in epidermis as well as pharynx (122). Cluster 23 also uniquely expresses several other genes that lack WISH data but may represent new epidermal markers including *ca6, cldn1, cxl34b.11, anxa2a, si:dkey-33c14.3, and mmp30*.

Cluster 26 is enriched for genes known to be expressed in the outermost layer of the epidermis called the peridermis including *krt4* (141) and anxa1c (125). Several genes enriched in this cluster (*dhrs13a.2, evpla, cldne, zgc:110333, zgc:111983*) have known expression in the periderm or other epidermal cells as well as the pharynx (122, 125, 142). Several other genes that lack available WISH data also showed strong expression in these cells including *icn2, zgc:153665, krt17, scel, and si:dkey-247k7.2*. Several genes were enriched in both clusters 23 and 26 including epidermal markers krtt1c19e (143), *cyt1* (124), and other genes like *cyt1l and spaca4l*. Taken together, these data indicate that clusters 23 and 26 represent cell types within the epidermis, with cluster 23 potentially representing basal cells and cluster 26 potentially representing peridermal cells. Considering that several of these genes are known to be expressed in epidermis as well as pharynx, it is possible some of these cells are located in the pharynx.

*Other cell types:* Our dataset identified several clusters that appear to be contaminating cell types not derived from the intestinal epithelium. For example, we annotate cluster 18 as leukocytes based on their expression of myeloid leukocyte markers including *spi1a, lcp1, mpeg1.1, and ncf1*. Cluster 19 we annotate as mesenchymal cells based on their expression of epithelial-mesenchymal transition genes like *twist1a and snai1a*, as well as vim and multiple collagen genes. Cluster 20 we annotate as exocrine pancreas based on expression of the pancreatic transcription factor *ptf1a* and other pancreatic markers *ela3l, pdia2, cpa1, prss1, amy2a* (122, 124, 144). Cluster 25 we annotate as red blood cells based on their unique expression of multiple hemoglobin genes. We were not able to confidently annotate cluster 24 due to a paucity of specific markers for which previous in situ hybridization data was available. Genes specific for cluster 24 include *cx30.3/gjb8* which is expressed in the *otic vesicle, skin, and swim bladder* (145, 146), and *si:dkey-96g2.1*. However, a known marker for the swim bladder epithelium mnx1/hb9 (147) was very low in our dataset, suggesting cluster 24 is not swim bladder. Several markers of cluster 24 were also shared with pharyngeal clusters 1, 7, and 10 (*clic2, tnfb, noxo1a, and CR762483.1*) suggesting some shared function between these cell types.

#### *In vitro* bile salt metabolism assay detects common bile salt modifications mediated by mammalian gut microbes

To test if complex gut microbiota or individual bacterial strains can modify bile salts, we developed an *in vitro* bile salt modification assay. Microbes of interest were first enriched in rich medium under aerobic or anaerobic conditions and were then incubated with known bile salts. Afterward, the metabolites from these cultures were extracted and subjected to LC/MS to examine potential modifications of the bile salts. To improve the detection sensitivity of the assay, we modified a previous extraction method used to extract cholesterol and other steroid hormones out of culture media for LC/MS analysis (*79*). Our extraction method was tested via extracting and quantifying 5αCS and TCA as well as ACA, a C_24_ bile acid analog of 5αCS, from modified TSB medium (Fig S3A). As expected, the internal standard D4-GCA was eluted at 5.62 min at 468.33 m/z; 5αCS was eluted at approximately 7.6 min at 531.3 m/z; TCA was eluted at approximately 6.2 min at 514.29 m/z; ACA was eluted at approximately 6.5 min at 407.29 m/z. These results confirmed that this method was appropriate for the extraction of 5αCS, TCA, and ACA from bacterial cultures. We next utilized several bacterial strains with known bile salt modification abilities as positive controls to validate the detection of bile salt modifications in our in vitro system. *Lactobacillus salivarius* JCM1046 encodes a bile salt hydrolase enzyme, which cleaves off the conjugated taurine in TCA, thus generating CA (*148*). *Clostridium scindens* ATCC 35704 is known to contain the bile acid 7α-dehydroxylation pathway, which can transform CA and/or ACA into secondary bile acids deoxycholic acid (DCA) or allo-DCA, respectively (*149*). Indeed, the predicted modified bile salts were recovered from the culture medium after incubation with the corresponding bacterial strains under anaerobic conditions (Fig S3B). Interestingly, we did not observe signs of 7α-dehydroxylation of 5αCS when supplementing this bile alcohol to *C. scindens* culture (Fig S3B). This suggests that 5αCS is likely resistant to such modification. Nonetheless, these data confirmed that our *in vitro* bile salt modification assay is appropriate for detecting modified metabolites of primary bile salts in cultures.

#### The Tg(-0.258fabp6-cfos:GFP)^rdu22^ is physically linked with cyp7a1 locus

To monitor Fxr activity in response to different levels of bile salts, we attempted to introduce the homozygous *cyp7a1^-16/-16^* mutation into the previously reported *Tg(-0.258fabp6- cfos:GFP)^rdu22^* transgenic line (*17*) by setting up breeding crosses between *cyp7a1^+/-16^* zebrafish and *cyp7a1^+/-16^; Tg(-0.258fabp6-cfos:GFP)* zebrafish. Following this cross, the progeny exhibited a normal Mendelian ratio with respect to the *cyp7a1* genotype and the wt: heterozygous: homozygous ratio was approximately 1:2:1 (Fig S1E). The segregation of *gfp* transgene also followed the Mendelian ratio, with only half of the progeny carrying the *gfp* (Fig S1E). Interestingly, we found that 152 out of 153 *gfp* positive offspring were either *cyp7a1*^+/+^ or *cyp7a1*^+/-^, whereas 158 out of 161 non-transgenic fish were either *cyp7a1*^+/-16^ or *cyp7a1*^-16/-16^. In fact, of all 314 progeny sampled, only one *cyp7a1*^-16/-16^; *Tg(-0.258fabp6-cfos:GFP)* fish and three *cyp7a1^+/+^* nontransgenic fish were identified. These genotypic frequencies suggested that the *gfp* transgene and *cyp7a1* gene were physically linked on chromosome 2 in the *Tg(- 0.258fabp6-cfos:GFP)* reporter line and therefore co-segregated following the cross. We therefore generated a new *Tg(-1.7fabp6:GFP)* reporter line and used the new line instead to examine Fxr activity upon bile salt deficiency (Fig 1D).

#### Validation of cyp7a1 mutant zebrafish

The *cyp7a1* mutant zebrafish were generated by targeting exon 2 that encodes the cytochrome P450 domain using CRISPR-Cas9. We identified two independent deletion alleles resulting in frameshift mutations in exon 2 of the *cyp7a1* gene, *cyp7a1*^-7/-7^ and *cyp7a1^-16/-16^*. Early mortality was observed in zebrafish homozygous for either of these *cyp7a1* alleles. Specifically, both the *cyp7a1*^-7/-7^ and *cyp7a1^-16/-16^* zebrafish showed exceptionally low survival rate past 1 month post-fertilization (1 out of >300 fish), consistent with the observations in *Cyp7a1* null mice (*150*). One allele, the larger *cyp7a1* deletion allele *cyp7a1^-16/-16^* (*rdu84*, designated as *cyp7a1^-/-^*) was selected for subsequent study. We validated this allele using two methods. First, we quantified the levels of bile salts in pooled *cyp7a1* wt versus homozygous mutant larvae. This revealed a significant reduction of the total bile salts in *cyp7a1* mutant zebrafish as compared to their wt counterparts, indicative of impaired bile salt synthesis in the *cyp7a1* mutant zebrafish. Second, we compared the mRNA levels of *cyp7a1* between larvae pools enriched for either *cyp7a1* wt or mutant zebrafish. Specifically, we crossed the *cyp7a1^+/-^* zebrafish with the *cyp7a1^+/-^; Tg(-0.258fabp6-cfos:GFP)* zebrafish. As mentioned above, since the *gfp* gene in the *Tg(-0.258fabp6-cfos:GFP)* reporter line is physically linked with the *cyp7a1* gene, the GFP positive progeny produced by this cross will be either *cyp7a1*^+/+^ or *cyp7a1*^+/-^ (designated as the wt enriched pool), whereas the GFP negative progeny will be either *cyp7a1*^+/-^ or *cyp7a1*^-/-^ (designated as the mutant enriched pool). qPCR analysis of the GFP negative and negative populations revealed that the mutant enriched pool (GFP negative) exhibited a strong reduction of *cyp7a1* transcript levels as compared to the wt enriched pool (GFP positive), suggesting efficient knockdown of the *cyp7a1* gene in the mutants. We also observed decreased levels of *fabp6* and *slc10a2* RNA in the mutant enriched pool, consistent with the prediction that Fxr activity is reduced upon bile salt deficiency in the *cyp7a1* mutant.

## Supplementary Materials and Methods

### Chemicals, reagents, and materials

Corning polypropylene conical tubes (15 mL and 50 mL) were purchased from MilliporeSigma (San Jose, CA). Organic solvents and other chemicals were purchased from Fisher Scientific (Hampton, NH). JT Baker 20 x 20 thin layer chromatography (TLC) plates were purchased from Thomas Scientific (Swedesboro, NJ). Solid phase extraction cartridges were purchased from Waters (Milford, MA). Allocholic acid was purchased from Cayman Chemicals (Ann Arbor, MI), and deuterated-glycocholic acid was purchased from Toronto Research Chemicals (Ontario, Canada). All other bile standards were purchased from MilliporeSigma. Other materials were purchased from either MilliporeSigma or Fisher Scientific. Since we determined that animal-based peptones contained detectable bile acids, vegetable peptone and soy peptone both from BD Biosciences (San Jose, CA) were used in culture medium. Other ingredients include glucose, sodium chloride, dipotassium phosphate, Tween 80, and cysteine obtained from Fisher Scientific.

### Total bile salt quantification

For whole larvae samples, 6 dpf larvae were euthanized and rinsed with GZM for three times to remove carryover of food or debris (30 larvae/replicate; 4-6 replicates/condition). The pooled larvae were then placed in 1 mL 1:1 chloroform: methanol and sonicated for 3 min with 2/1 ON/OFF cycle and 70% amplitude using a QSONICA Q700-MPX-110 cup horn sonicator. The homogenate of each replicate was spun at room temperature at 20000 g for 30 min. The supernatant was collected, air-dried in a fume hood overnight, and subjected to bile salt quantification using the enzyme-cycling method based bile acid detection kit (Diazyme, DZ042A) as described previously (*14*).

### In vitro *bile salt modification assay*

Modified tryptic soy broth (TSB) at pH= 7.0 was prepared aerobically and anaerobically for experimental use with the composition as follows: vegetable peptone (17.0 g/L), soy peptone (3.0 g/L), glucose (2.5 g/L), sodium chloride (0.5 g/L), dipotassium phosphate (2.5 g/L), and cysteine (1.0 g/L). The substrate testing medium was generated by supplemented freshly made modified TSB medium with 50 µ M methanol vehicle, 5αCS, TCA, or ACA, respectively. Two microbes with known bile salt conversion activities were selected as reference controls for bile acid and bile alcohol conversion: *Lactobacillus salivarius* JCM1046 for the bile salt hydrolase activity (*151*), and *Clostridium scindens* ATCC 35704 for 7α-dehydroxylation activity (*149*). Bacterial reference control tubes were also inoculated (1:10 dilution) in modified TSB medium with the selected strains of *L. salivarius* and *C. scindens* containing 50 µ M corresponding bile salt substrates and cultured at 37 °C for 48 h. For *L. salivarius*, 1% Tween 80 was supplemented to media for growth enrichment (*152*).

---

**Supplementary Figure 1.**
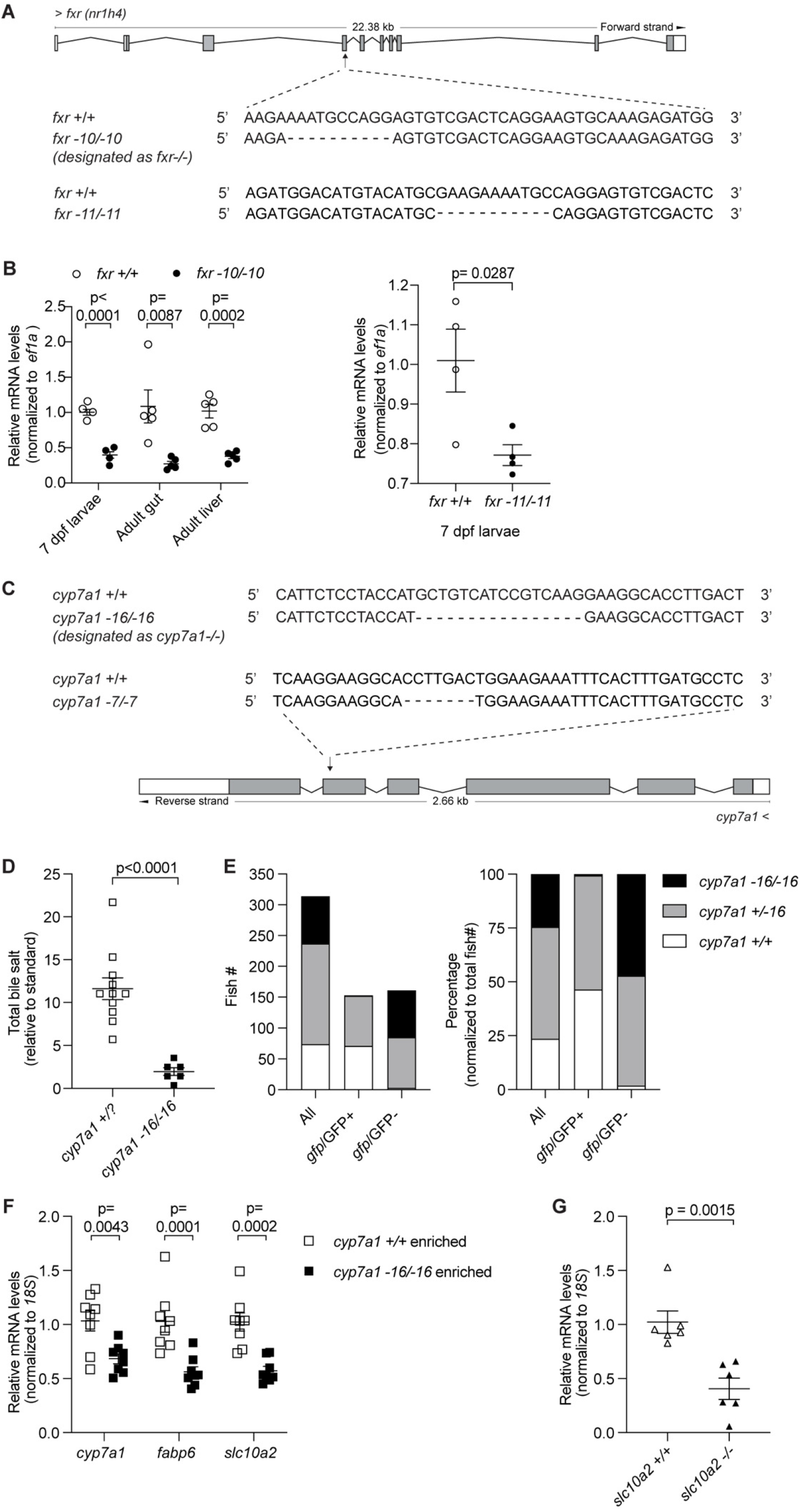
Zebrafish mutants with disrupted bile salt signaling pathways are validated by qRT-PCR and chemical assay. **A.** Schematic representation of the zebrafish *fxr* locus along with the wt and the indel sequences from two independently identified alleles. Nucleotide changes are listed as dashes. **B.** qRT-PCR analysis comparing the expression of *fxr* between *fxr^+/+^* and *fxr^-10/-10^* (left) or *fxr^-11/-11^*(right) zebrafish at different developmental stages and in different tissues. The results are represented as relative expression levels that were normalized to *ef1a* (Mean±SEM). **C.** Schematic representation of the zebrafish *cyp7a1* locus along with the wt and indel sequences from two independent alleles. Nucleotide changes are listed as dashes. **D.** Total bile salt levels in whole larvae of 6 dpf *cyp7a1^+/?^* (WT or heterozygous) and *cyp7a1^-16/-16^* (designated as *cyp7a1^-/-^*, homozygous) zebrafish. E. Distribution of cyp7a1 and gfp genotypes in progeny from cross between *cyp7a1^+/-16^* and *cyp7a1^+/-16^*; Tg(-0.258fabp6-cfos:GFP) zebrafish. Raw fish counts of the respective genotype are shown on the left. The percentage of each genotype relative to the total fish counts is shown on the right. **F.** qPCR analysis comparing the expression of the direct Fxr target genes in 6 dpf *cyp7a1^+/+^* enriched larvae and *cyp7a1^-16/-16^* enriched larvae. G. qPCR analysis comparing expression of slc10a2 in 6 dpf *slc10a2^+/+^* and *slc10a2^sa2486/sa2486^* larvae. The results in (B, F, G) are represented as relative expression levels normalized to *18S* (Mean±SEM). Statistical significance in (**B, D, G**) was calculated by unpaired t-test and in (**F**) was calculated by two-way ANOVA with Turkey’s multiple comparisons test. Shown are representative data from at least two independent experiments.

**Supplementary Figure 2.**
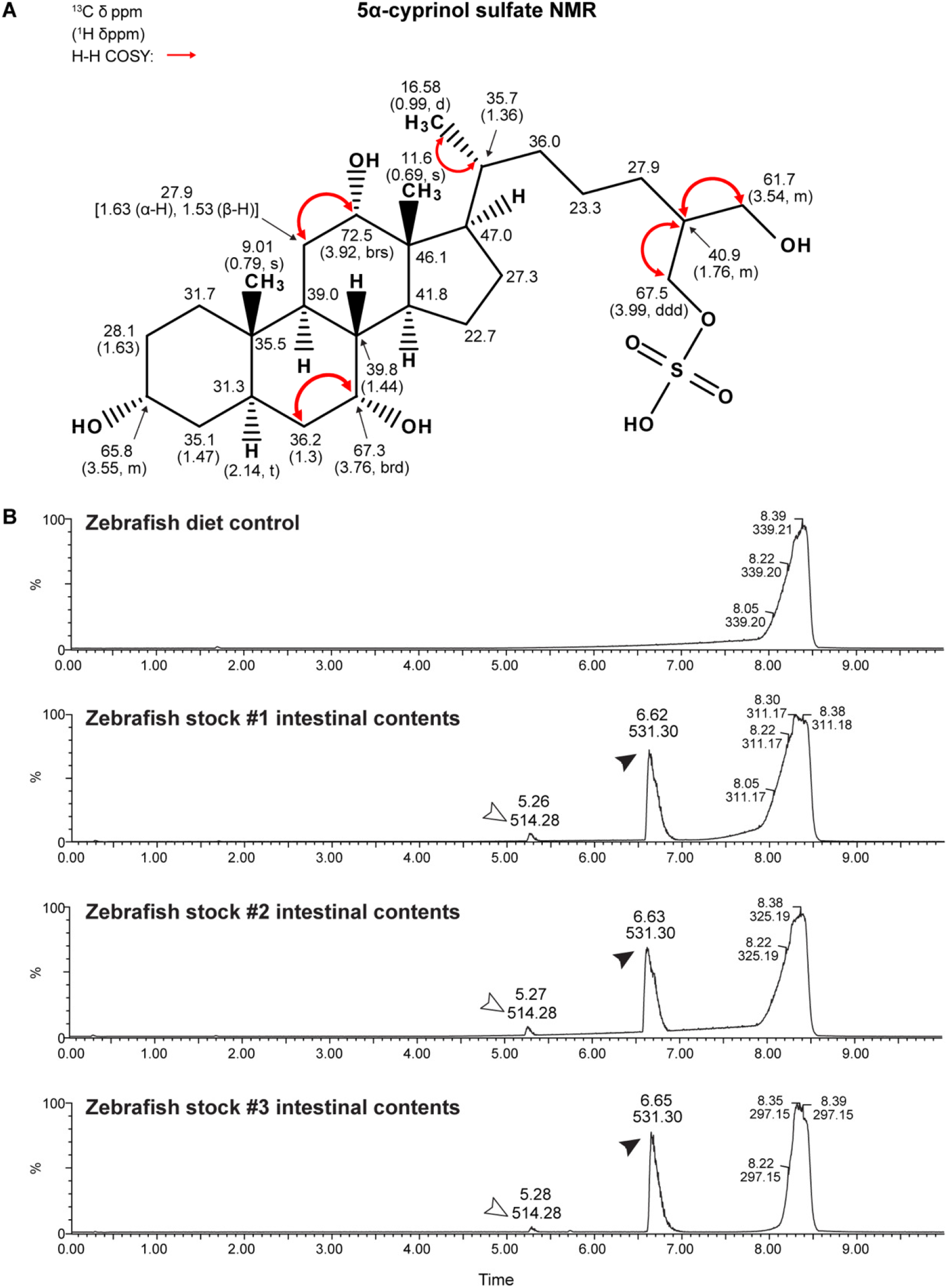
The major zebrafish biliary bile alcohol species is confirmed by NMR as 5α-cyprinol sulfate and is detected in the zebrafish intestinal contents. **A.** Proton and Carbon-13 NMR chemical shifts of 5α-cyprinol sulfate isolated from zebrafish bile. The carbon-13 chemical shifts were determined relative to the CD3OD resonance and converted to the tetramethylsilane (TMS) scale using δ (CD3OD) = 49.3 ppm. Values in parentheses refer to proton chemical shifts (δ ppm from TMS) and signal multiplicity: singlet, s; multiplet, m; doublet, d; triplet, t; brd, broad doublet; double doublet, dd. Arrows represents observed proton heteronuclear multiple bond correlations (HMBC). **B.** LC/MS chromatograms of metabolites extracted from the diet or intestinal contents of adult zebrafish from three randomly selected fish stocks. The arrowheads indicate the major zebrafish bile salts: black: 5αCS; white: TCA.

**Supplementary Figure 3.**
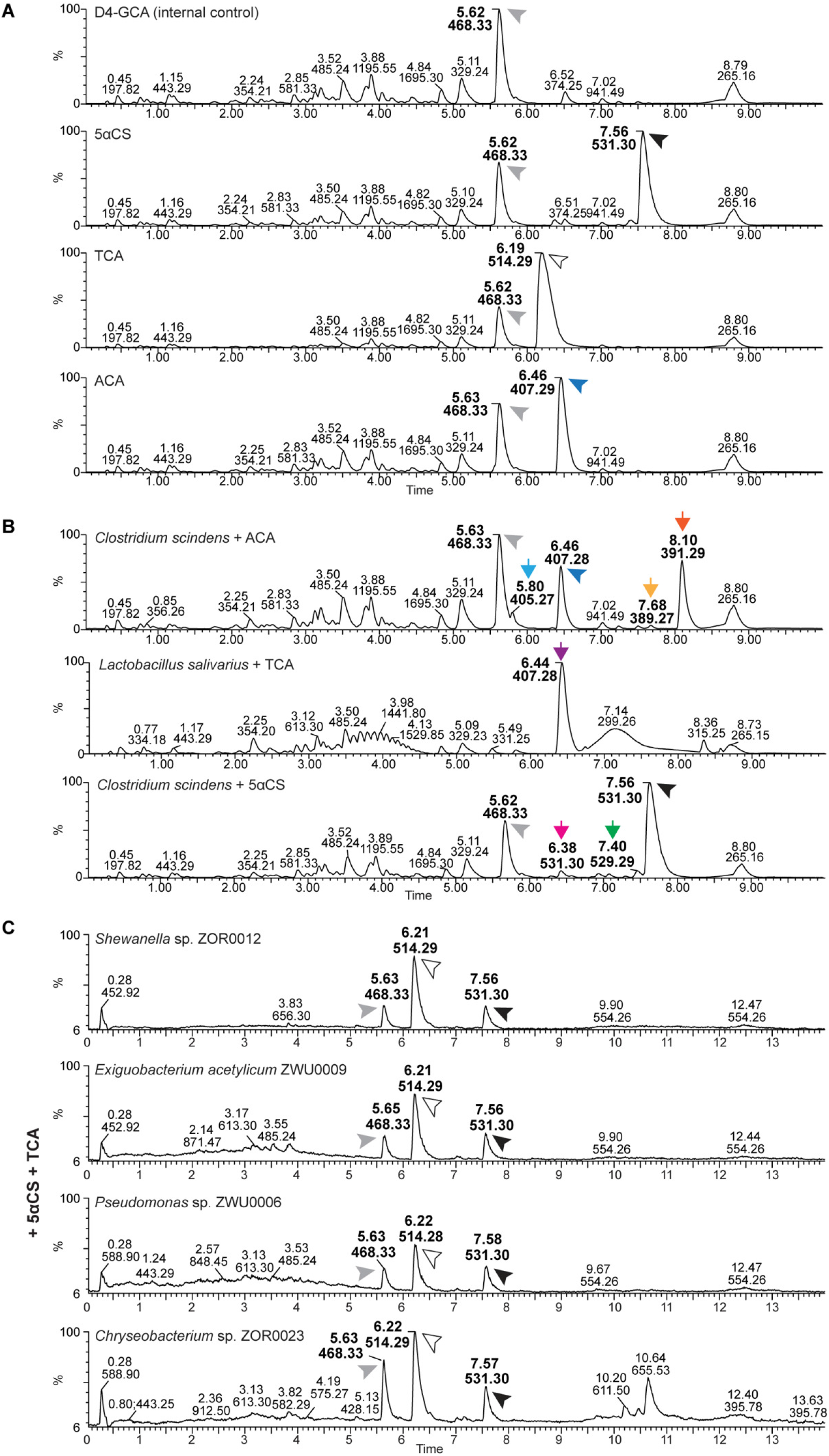
*In vitro* bile salt metabolism assay reveals specific modifications of bile salts by mammalian or zebrafish gut microbes. **A.** LC/MS chromatographs of metabolites extracted from medium incubated with internal standard D4-GCA, 5αCS, TCA, and ACA for 24 hours. The arrowheads indicate the supplemented substrates: grey: internal standard D4-GCA; black: 5αCS; white: TCA; blue: ACA. **B.** LC/MS chromatographs of bile salt metabolites extracted from enriched bacterial monocultures supplemented with 50 μM known bile salts under anaerobic conditions. The bacteria and bile salt combinations are as follows: top: *Clostridium scindens* ATCC 35704 with ACA; middle: *Lactobacillus salivarius* JCM1046 with TCA; bottom: *Clostridium scindens* ATCC 35704 with 5αCS. The arrowheads indicate the supplemented substrates: grey: internal standard D4-GCA; black: 5αCS; white: TCA; blue: ACA. The arrows indicate bile salt metabolites resulted from microbial modification of the supplemented bile salts: light blue: dehydrogenated ACA; dark orange: allo-DCA; light orange: dehydrogenated allo-DCA; purple: CA; green: dehydrogenated 5αCS; magenta: epimerized 5αCS. No other modifications of bile salts were observed. C. LC/MS chromatographs of bile salt metabolites extracted from enriched monocultures of zebrafish intestinal microbes supplemented with 50 μM 5αCS and TCA under aerobic conditions. The bacteria were chosen to represent the major bacterial taxa in zebrafish gut and are listed as follows (from top to bottom): *Shewanella* sp. ZOR0012 (Proteobacteria), *Exiguobacterium acetylicum* ZWU0009 (Firmicutes), *Pseudomonas sp.* ZWU0006 (Proteobacteria), *Chryseobacterium sp.* ZOR0023 (Bacteroidetes). The arrowheads indicate the supplemented substrates: grey: internal standard D4-GCA; black: 5αCS; white: TCA. No modification of 5αCS and TCA by these bacteria was observed.

**Supplementary Figure 4.**
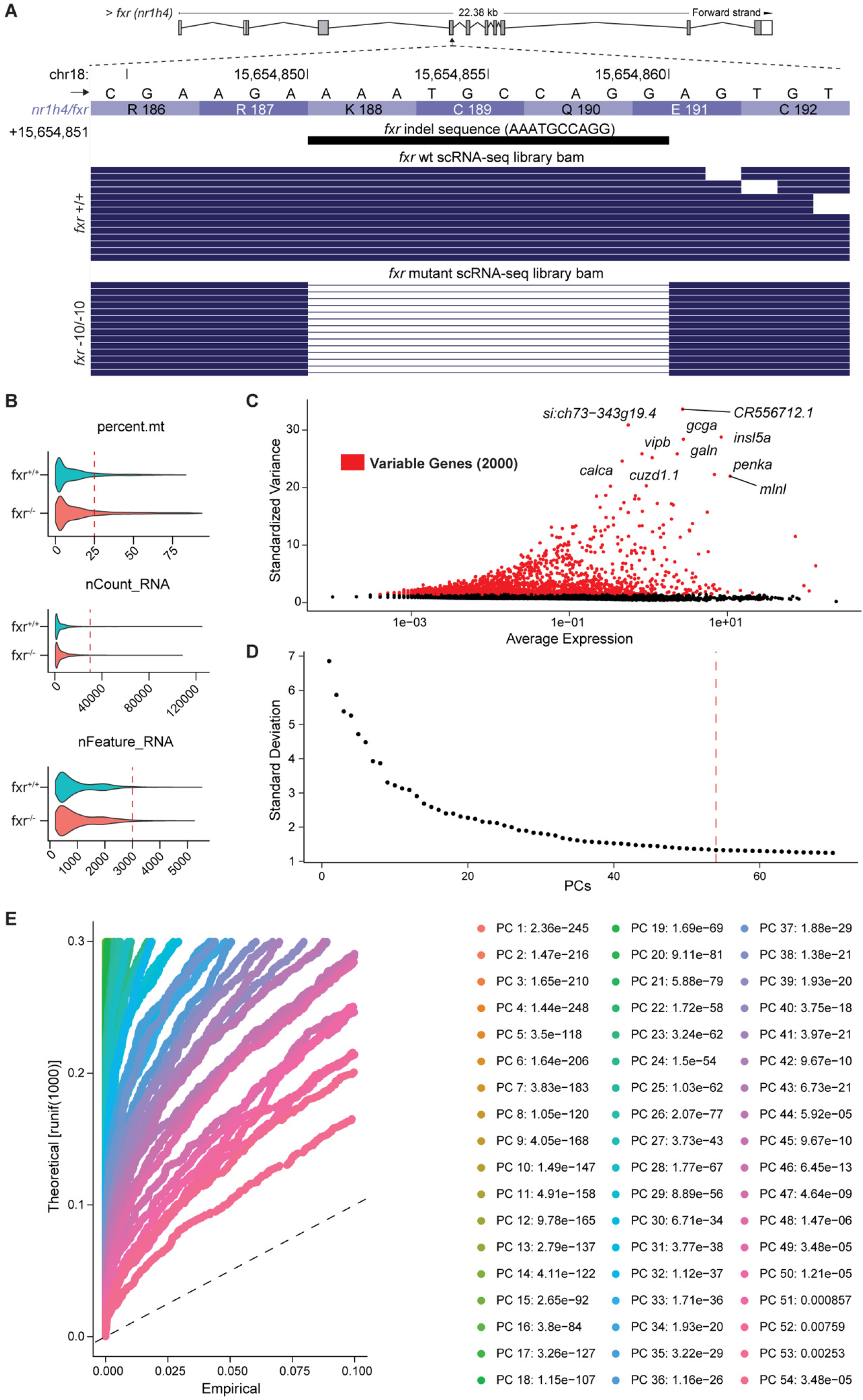
ScRNA-seq data are processed for quality control prior to clustering. **A.** Diagram representing a stack of randomly selected reads spanning the fxr exon 4 from the scRNA-seq, aligned to the exact indel sequence of *fxr^-/-^*. **B.** Number of counts per cell, features (or genes) per cell and percent mitochondrial transcripts for both *fxr^+/+^* (blue) and *fxr^-/-^* (pink) libraries. Dash lines in each panel represent the QC threshold cutoff used to select cells for downstream analyses. **C.** Highly variable genes (in red) determined by the average expression and vst method. **D.** ElbowPlot (100 PCs included, red dashed line at 54) and **E.** Jackstraw analysis (54 significant PCs included) employed to determine the maximum number of PCs that significantly contributed to variation in the integrated dataset.

**Supplementary Figure 5.**
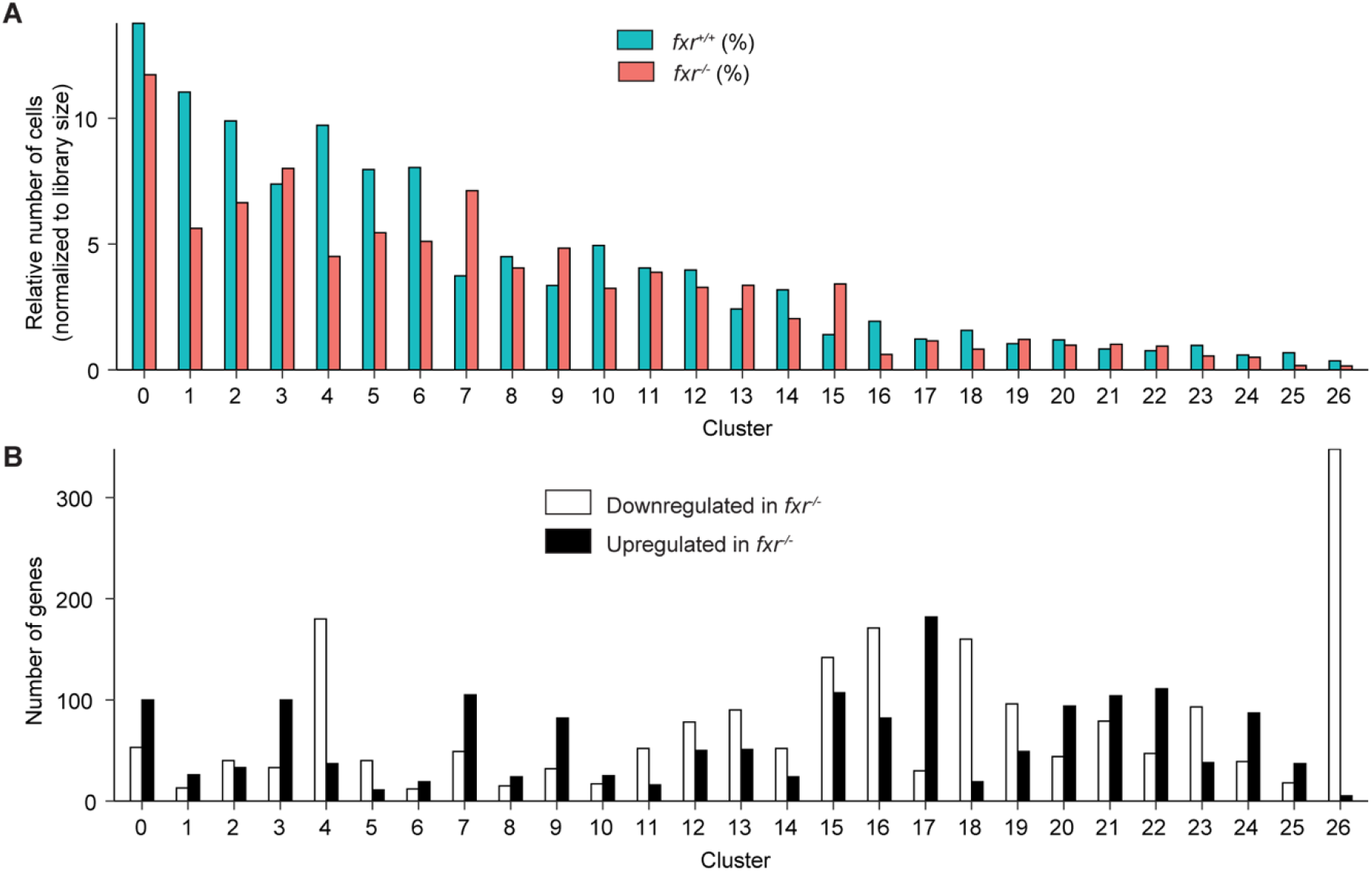
Fxr mutation displays broad impacts on relative abundance and gene expression in multiple cell types of zebrafish intestinal epithelial cells. **A.** Relative abundance of the *fxr^+/+^* and *fxr^-/-^* cells in each cluster. The relative abundance was calculated by dividing the raw cell abundances of a cluster by the total number of cells per genotype and represented as a percentage. **B.** Total number of genes upregulated or downregulated in *fxr^-/-^* samples by cluster.

**Supplementary Figure 6.**
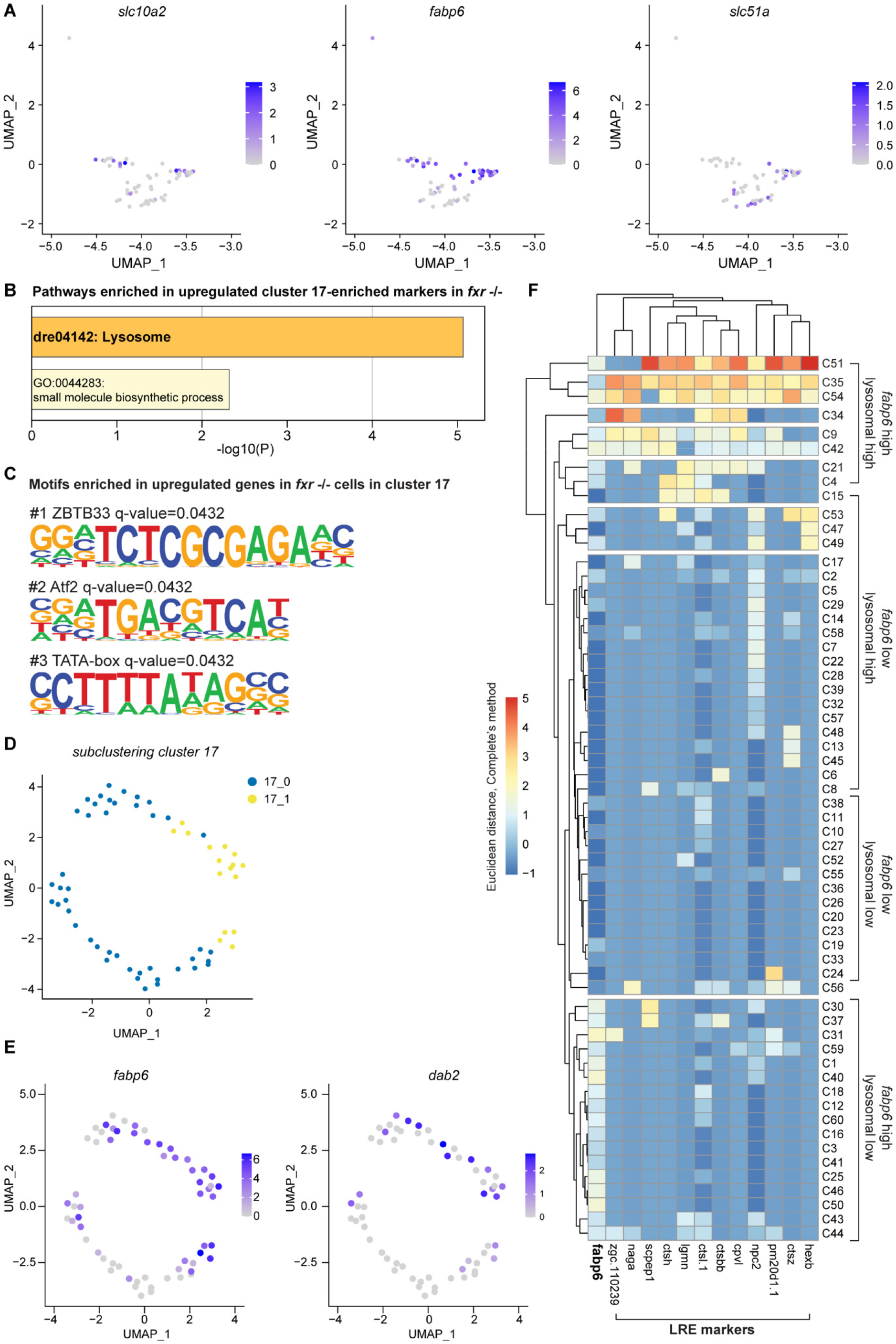
Single-cell RNA sequencing highlights transcriptomic heterogeneity in zebrafish ileal epithelial cells. **A.** UMAP plots revealing expression and distribution of genes related to bile salt transport in cluster 17 cells of the *fxr* wt zebrafish. **B.** Non-redundant enrichment terms identified by Metascape upon analysis of upregulated cluster 17-enriched markers of *fxr^-/-^* cells. The colors of enriched terms are scaled to represent statistical significance. **C.** The top 3 HOMER-identified enriched motifs in genes that were up regulated in the *fxr^-/-^* cells relative to *fxr^+/+^* cells in cluster 17. Shown are the position weight matrices (PWMs) of the enriched nucleotide sequences. The TF family that most closely matches the motif is indicated above the PWM. **D.** UMAP plot revealing two subclusters (17_0 and 17_1) of wt cells in cluster 17. **E.** FeaturePlots of ileocyte marker *fabp6* and LRE marker *dab2* showing concurrent expression in cluster 17 cells. **F.** Hierarchical clustering of *fxr^+/+^* cluster 17 cells based on expression of bile absorption marker and lysosomal markers. Each row represents an individual cluster 17 cell and each column represents the expression of the indicated gene. Cells are manually assigned into four groups based on their expression profiles: *fabp6* high/lysosomal marker high; *fabp6* low/lysosomal marker high; *fabp6* low/lysosomal marker low; *fabp6* high/lysosomal marker low.

**Supplementary Figure 7.**
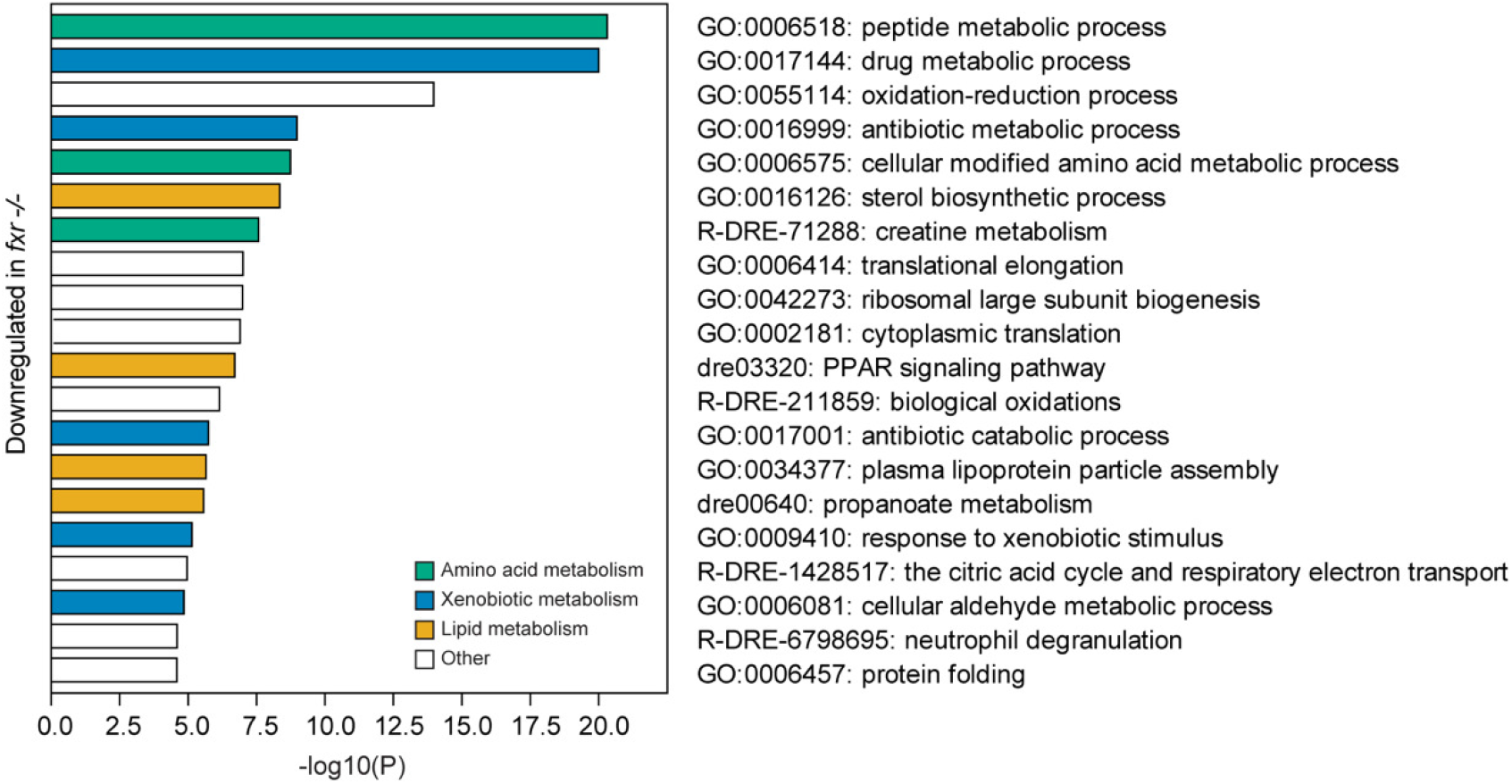
Metabolism pathways are enriched within genes downregulated in cluster 4 cells upon *fxr* mutation. Top 20 non-redundant enrichment terms identified by Metascape upon analysis of genes that were down regulated in the *fxr^-/-^* cells relative to *fxr^+/+^* cells in cluster 4. Enriched terms related to metabolism are color coded as follows: orange: lipid metabolism; green: amino acid metabolism; blue: xenobiotic metabolism; white: others.

**Supplementary Figure 8.**
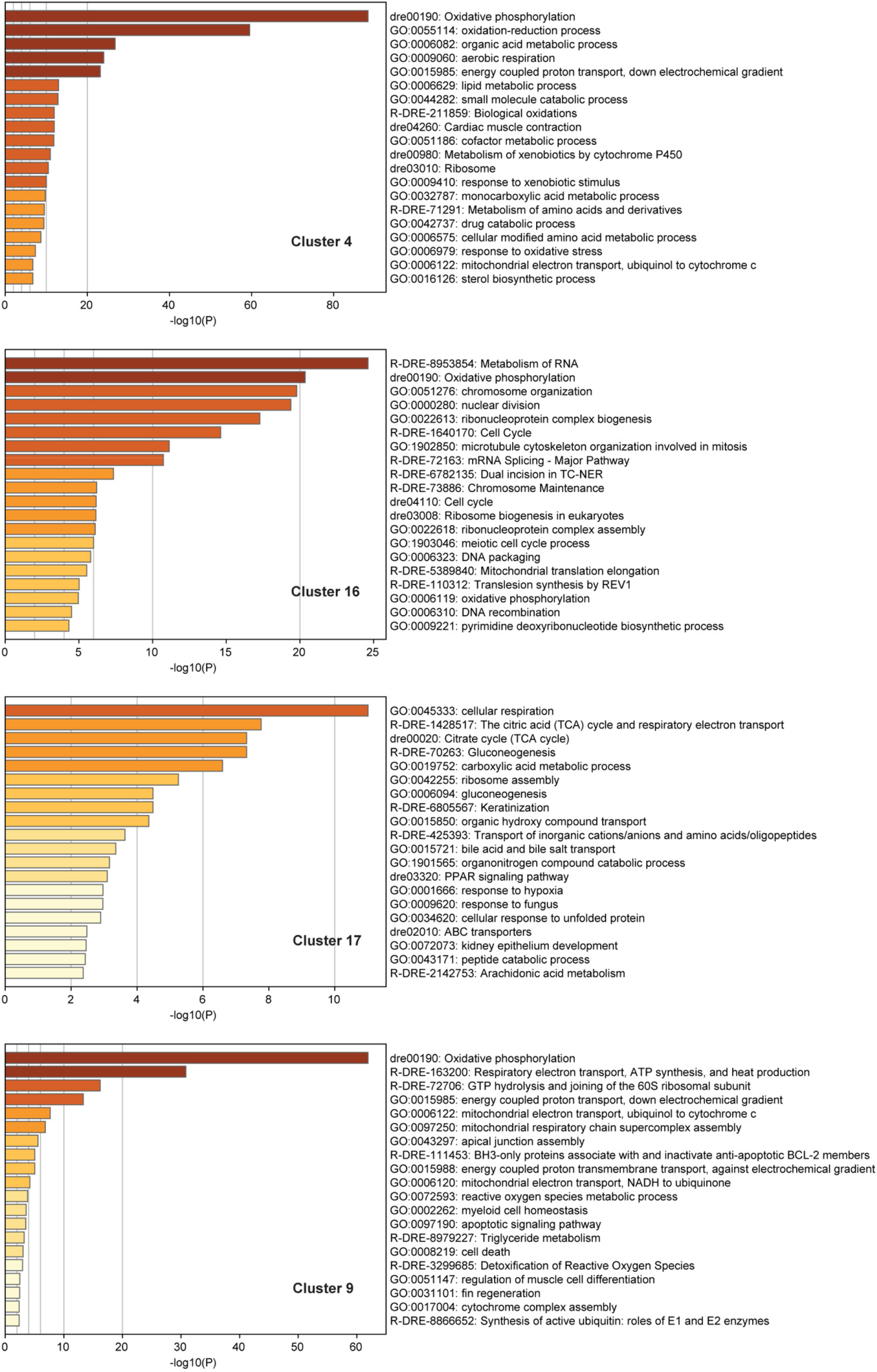
Functional categorization analysis reveals enriched pathways in each enterocyte cluster. Top 20 non-redundant enrichment terms identified by Metascape upon analysis of marker genes of each enterocyte cluster defined by our scRNA-seq analysis. The colors of enriched terms are scaled to represent statistical significance. Shown from top to bottom are functional enrichment analysis of cluster 4, 16, 17, and 9.

**Supplementary Table 1.**
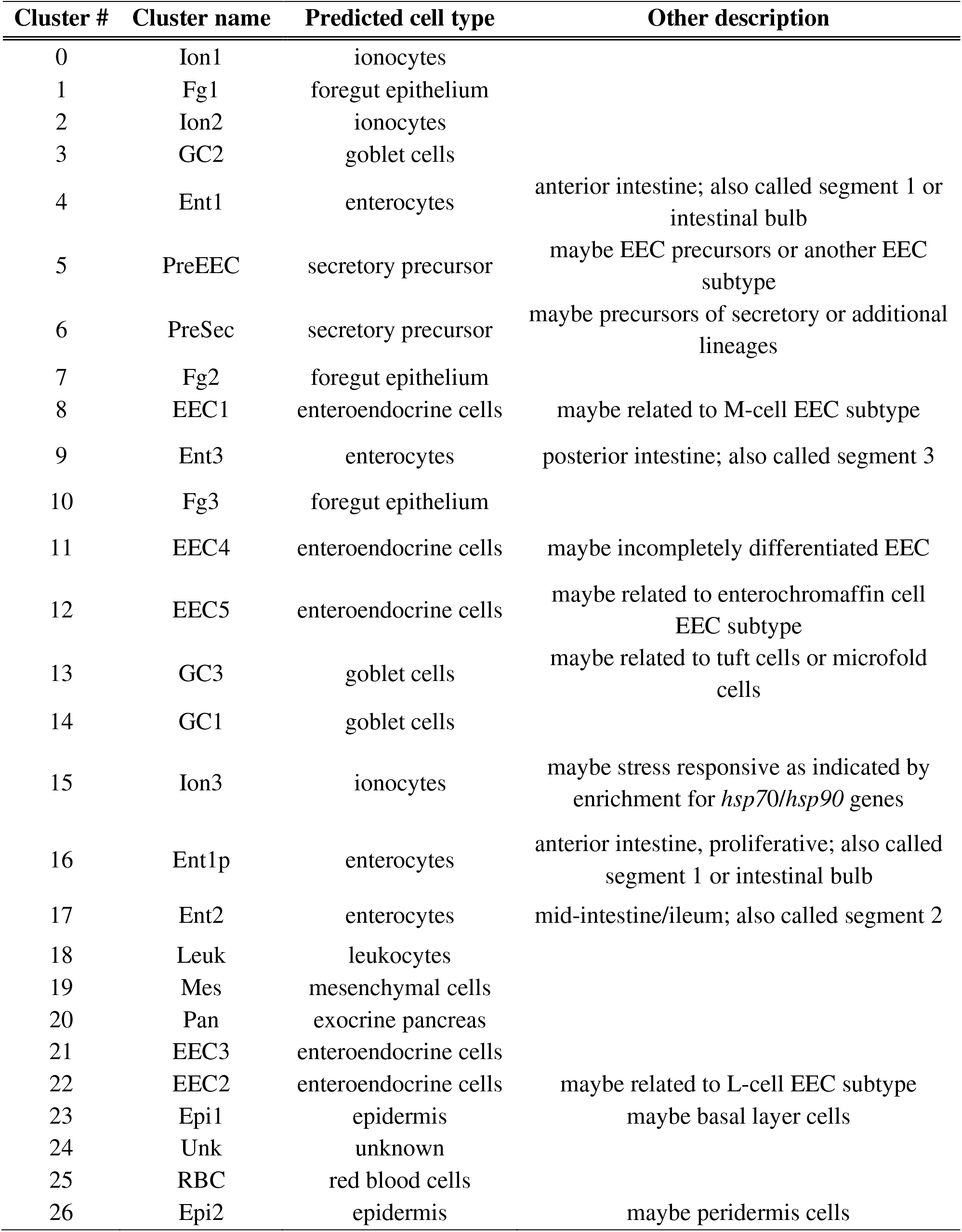
Annotation of cell clusters based on single-cell RNA-seq analysis of wild-type zebrafish intestinal epithelial cells.

**Supplementary Table 2.**
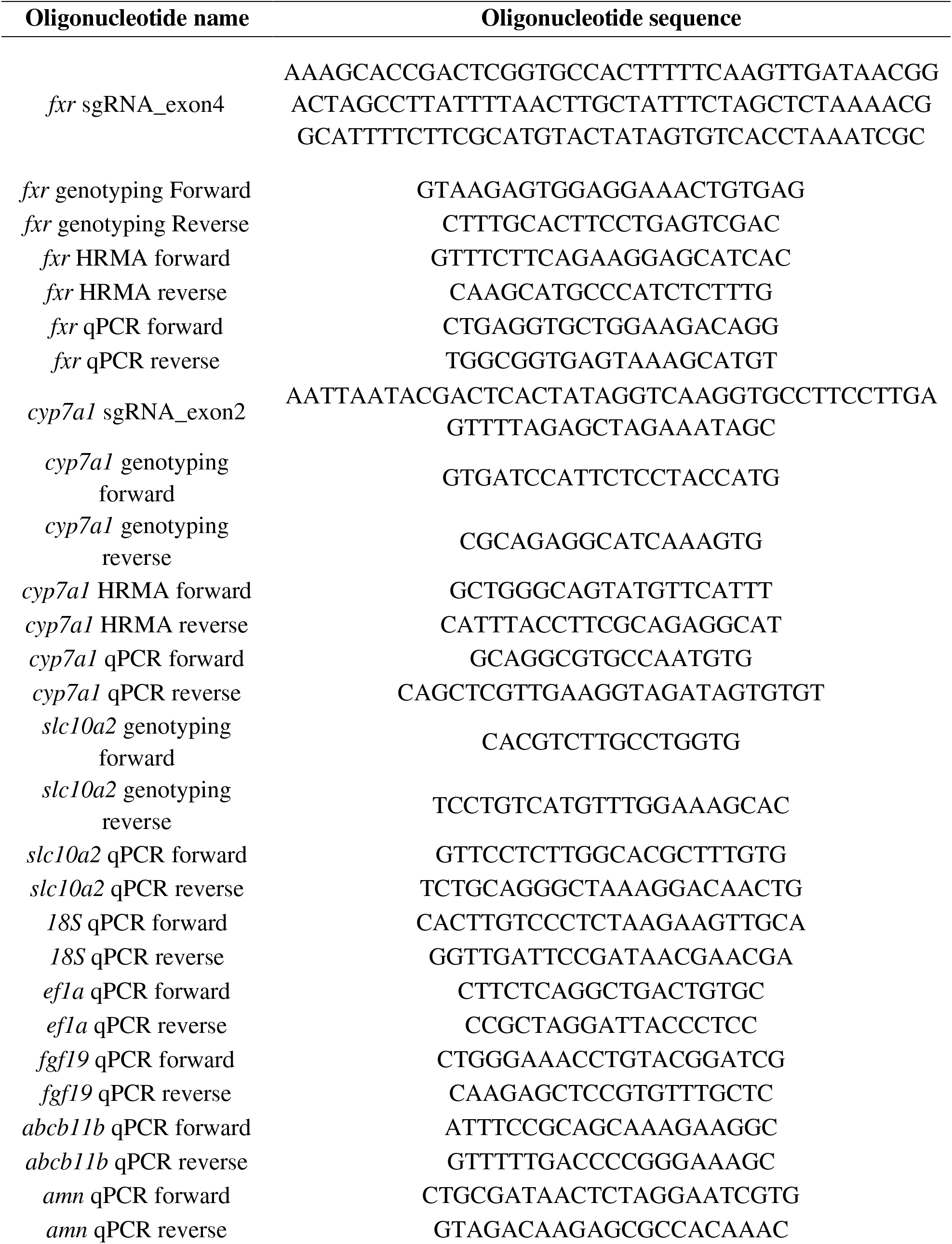
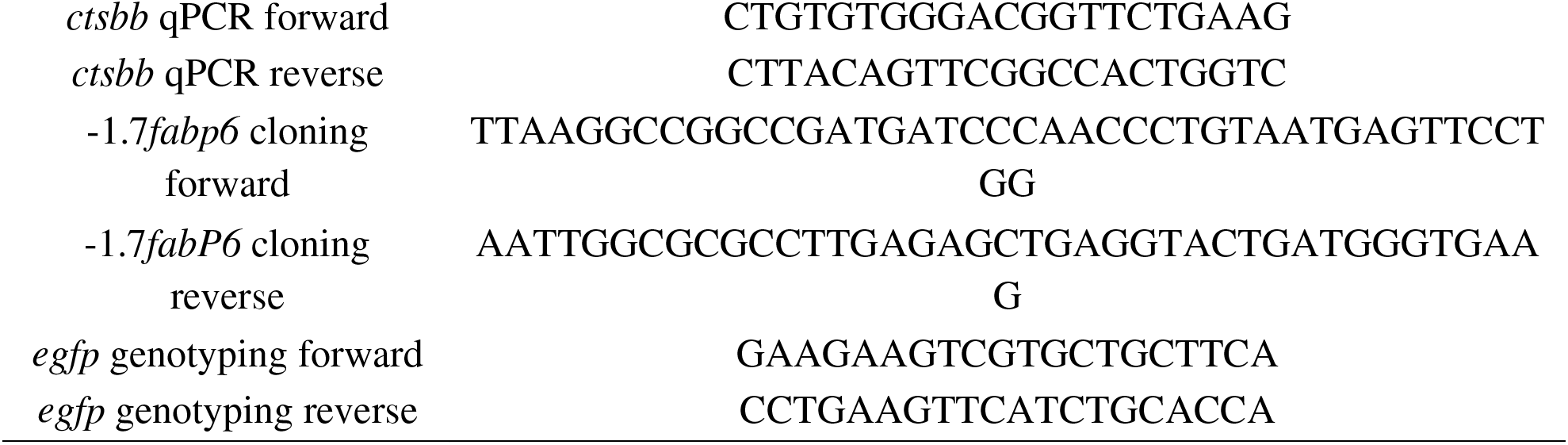
Oligonucleotides used in this study

## Supplementary data files S1-S5

Dataset 1. Average expression of genes in individual cell clusters identified by scRNA-seq analysis of wild-type zebrafish intestinal epithelial cells

Dataset 2. Cluster markers of individual cell clusters identified by scRNA-seq analysis of wild-type zebrafish intestinal epithelial cells

Dataset 3. Cluster-enriched markers of individual clusters identified by scRNA-seq analysis of wild-type zebrafish intestinal epithelial cells

Dataset 4. Differentially expressed genes between fxr wild-type and mutant cells in individual cluster clusters identified by scRNA-seq analysis of wild-type zebrafish intestinal epithelial cells

Dataset 5. Comparisons of Fxr regulon in larval zebrafish intestine and adult mouse intestine

